# Elevated pyramidal cell firing orchestrates arteriolar vasoconstriction through COX-2-derived prostaglandin E2 signaling

**DOI:** 10.1101/2024.03.27.586974

**Authors:** Benjamin Le Gac, Marine Tournissac, Esther Belzic, Sandrine Picaud, Isabelle Dusart, Hédi Soula, Dongdong Li, Serge Charpak, Bruno Cauli

## Abstract

Neurovascular coupling, linking neuronal activity to cerebral blood flow, is essential for brain function and underpins functional brain imaging. Whereas mechanisms involved in vasodilation are well-documented, those controlling vasoconstriction remain overlooked. This study unravels the mechanisms by which pyramidal cells elicit arteriole vasoconstriction. Using patch-clamp recording, vascular and Ca^2+^ imaging in mouse cortical slices, we show that strong optogenetic activation of layer II/III pyramidal cells induces vasoconstriction, correlating with firing frequency and somatic Ca^2+^ increase. *Ex vivo* and *in vivo* pharmacological investigations indicate that this vasoconstriction predominantly recruits prostaglandin E2 through the cyclooxygenase-2 pathway, and activation of EP1 and EP3 receptors. We also present evidence that specific interneurons releasing neuropeptide Y, and astrocytes, through 20-hydroxyeicosatetraenoic acid, contribute to this process. By revealing the mechanisms by which pyramidal cells lead to vasoconstriction, our findings shed light on the complex regulation of neurovascular coupling.

**Significance statement:** Cerebral blood flow is tightly controlled by neuronal activity, a process termed neurovascular coupling which serves as the physiological basis for functional brain imaging widely used to map neuronal activity in health and diseases. While the prevailing view links increased neuronal activity with enhanced blood perfusion, our data suggest that elevated neuronal activity can also reduce cerebral blood flow. By optically controlling the activity of pyramidal cells, we demonstrate that these excitatory neurons induce vasoconstriction when their action potential firing is increased by releasing glutamate and lipid messengers. These findings update the interpretation of functional brain imaging signals and help to better understand the etiopathogenesis of epilepsy and Alzheimer’s disease, in which hyperactivity, hypoperfusion and cognitive deficits overlap.

## Introduction

The brain critically depends on the uninterrupted blood supply provided by a dense vasculature (Schmid et al., 2019). Cerebral blood flow (CBF) is locally and temporally controlled by neuronal activity, by an essential process called neurovascular coupling (NVC), and is impaired in early stages of numerous neurological disorders (Iadecola, 2017). NVC also serves as the physiological basis for functional brain imaging widely used to map neuronal activity. Neuronal activity increases CBF within seconds (Iadecola, 2017). In the cerebral cortex, the hyperemic response linked to neural activity is supported by dynamically controlled vasodilation that is spatially and temporally constrained by vasoconstriction in a second phase (Devor et al., 2007). Conversely, vasoconstriction and decreased CBF usually correlate with reduced neuronal activity (Devor et al., 2007; Shmuel et al., 2002).

Mounting evidence indicates that the positive correlation between neuronal activity and CBF is not always maintained under physiological conditions: i) robust sensory-evoked vasodilation can occur in the absence of substantial neuronal response (O’Herron et al., 2016), ii) conversely, pronounced neuronal activity is not systematically associated with increased hemodynamics (Ma et al., 2016), iii) CBF is decreased in several cortical areas despite local increase in neuronal activity (Devor et al., 2008), and iv) optogenetic simulation of inhibitory GABAergic interneurons results in vasodilation (Uhlirova et al., 2016). Furthermore, in pathological conditions with intense neuronal activity such as epileptic seizures, a sustained hypoperfusion induced by vasoconstriction is observed (Farrell et al., 2016; Tran et al., 2020).

NVC is achieved by the synthesis and release of vasoactive messengers within the neurovascular unit (Iadecola, 2017), which act on the contractility of mural cells (smooth muscle cells and pericytes) to control vessel caliber and CBF along the vascular tree (Rungta et al., 2018). Pial and penetrating arterioles, which have a higher density of contractile mural cells and control their diameter faster than capillaries (Hartmann et al., 2021; Hill et al., 2015; Rungta et al., 2021, 2018), play a key role in regulating CBF.

Different experimental approaches, each with their advantages and limitations, have allowed the identification of several mediators of NVC (Grutzendler and Nedergaard, 2019; Iadecola and Nedergaard, 2007). *Ex vivo* brain slices provide a well-controlled environment, ideal for pharmacological investigations to dissect the underlying mechanisms. However, they lack connectivity and blood flow which provides both vascular tone and natural oxygenation and therefore require *in vivo* validation. Conversely, pharmacological studies are more challenging with i*n vivo* preparations. Awake animals allow physiologically relevant context with largely undisturbed network and neuromodulatory activity. However, this preparation is subject to brain state changes which may affect network activity, metabolism, and vascular physiology (Grutzendler and Nedergaard, 2019), potentially complexifying the analysis of specific mechanisms of NVC. Although chronic anesthetized animals have reduced network and neuromodulation activity, the NVC response is only slowed (Rungta et al., 2021), providing a valuable model for validating *ex vivo* observations.

Messengers of vasodilation released by excitatory neurons, GABAergic interneurons, astrocytes, or endothelial cells, include nitric oxide, K^+^, arachidonic acid derivatives such as prostaglandin E2 (PGE2) (Iadecola, 2017), or more recently glutamate (Zhang et al., 2024). Despite its physio pathological importance, vasoconstriction is less understood with fewer cell types and vasoactive messengers that have been identified. It is now generally accepted that GABAergic interneurons are key players in vasoconstriction by releasing neuropeptide Y (NPY) (Cauli et al., 2004; Uhlirova et al., 2016). Under certain conditions, astrocytes can also induce vasoconstriction via 20-hydroxyeicosatetraenoic acid (20-HETE) (Mulligan and MacVicar, 2004) or high K^+^ concentration (Girouard et al., 2010). However, the involvement of pyramidal cells in vasoconstriction has been overlooked.

PGE2 has emerged as a bimodal messenger of NVC, similar to K^+^ (Girouard et al., 2010) and glutamate (Zhang et al., 2024), that can induce either vasodilation (Gordon et al., 2008; Lacroix et al., 2015; Lecrux et al., 2011; Mishra et al., 2016) or vasoconstriction (Dabertrand et al., 2013; Rosehart et al., 2021) depending on its concentration and/or site of action along the vascular tree. Under physiological conditions, PGE2 is produced during NVC by either astrocytes (Mishra et al., 2016) or pyramidal cells (Lacroix et al., 2015) via the rate-limiting synthesizing enzymes cyclooxygenase-1 (COX-1) or −2 (COX-2), respectively. Since COX-2-expressing pyramidal cells can release glutamate and PGE2, both of which induce vasoconstriction at high concentrations (Dabertrand et al., 2013; Rosehart et al., 2021; Zhang et al., 2024), pyramidal cells may be responsible for vasoconstriction when their spiking activity is high.

To test this hypothesis, we used *ex vivo* and *in vivo* approaches in combination with optogenetics to precisely control pyramidal cell firing in the mouse barrel cortex while monitoring the resulting arteriolar response. We found that pyramidal cells induce vasoconstriction at high stimulation frequency and about half of them express all the transcripts required for a cell autonomous synthesis of the vasoconstrictor messengers PGE2 and prostaglandin F2α (PGF2α). Pharmacological investigations revealed that this neurogenic vasoconstriction depends on COX-2-derived PGE2 via the direct activation of vascular EP1 and EP3 receptors. It also involves the recruitment of intermediary NPY interneurons acting on the Y1 receptor, and, to a lesser extent astrocytes, via 20-HETE and COX-1-derived PGE2. Thus, our study reveals the mechanisms by which high frequency pyramidal cell firing leads to vasoconstriction.

## Results

### Pyramidal cells induce vasoconstriction at high firing frequency

To determine if pyramidal cells action potential (AP) firing can induce vasoconstriction in a frequency-dependent manner, we used optogenetics to induce AP firing while monitoring the resulting vascular response in cortical slices. We used Emx1-cre;Ai32 transgenic mice expressing the H134R variant of channelrhodopsin-2 (ChR2) in the cortical glutamatergic neurons (Gorski et al., 2002), conferring robust pyramidal cell photoexcitability (Madisen et al., 2012). Wide-field photostimulation of cortical slices was achieved in layers I to III (Supplementary Fig. 1a) using 10-second trains of 5 ms light pulses (see Methods) delivered at five different frequencies (1, 2, 5, 10 and 20 Hz, Fig. 1a).

**Figure 1:**
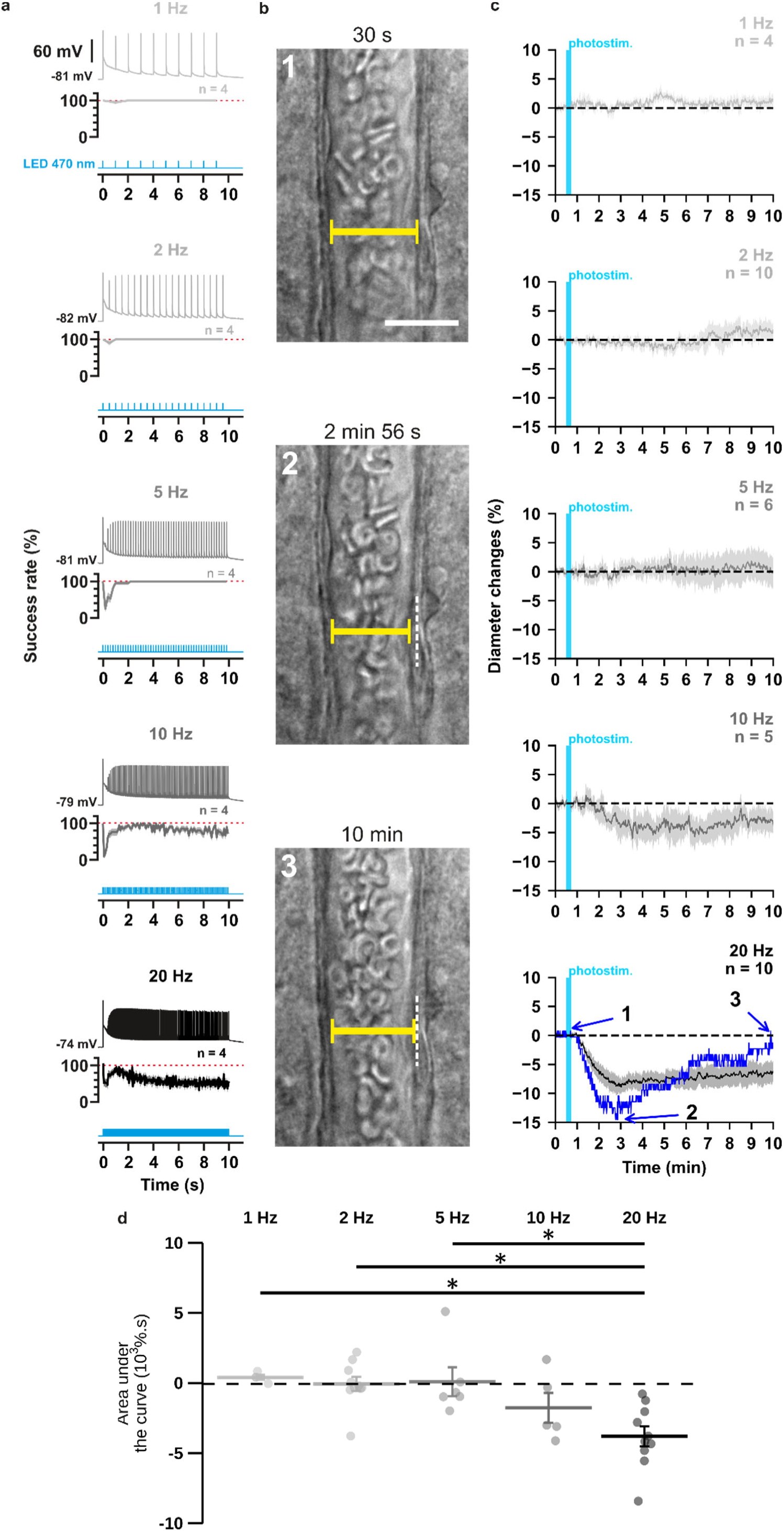
The occurrence and strength of vasoconstriction depends on the photostimulation frequency of pyramidal cells. **(a)** Representative examples of the voltage responses of a layer II-III pyramidal cell (upper traces light grey to black traces) induced by photostimulations (470 nm, 10 s train, 5 ms pulses) delivered at 1, 2, 5, 10 and 20 Hz (cyan lower traces) and mean spike success rate (middle trace, n= 4 cells from 3 mice). The SEMs envelope the mean traces. The red dashed lines represent a spike success rate of 100%. **(b)** Representative example showing IR-DGC pictures of a layer I penetrating arteriole (1) before a 20 Hz photostimulation, (2) at the maximal diameter decrease, and (3) after 10 minutes of recording. Pial surface is upward. Yellow calipers represent the measured diameters. White dashed lines indicate the initial position of the vessel wall. Scale bar: 25 µm. **(c)** Kinetics of arteriolar diameter changes induced by photostimulation (vertical cyan bars) at 1 Hz (n= 4 arterioles from 3 mice), 2 Hz (n= 10 arterioles from 8 mice), 5 Hz (n= 6 arterioles from 6 mice), 10 Hz (n= 5 arterioles from 5 mice) and 20 Hz (n= 10 arteriole from 9 mice). The SEMs envelope the mean traces. The blue trace represents the kinetics of the diameter changes of the arteriole shown in (b). **(d)** Effects of the different photostimulation frequencies on AUC of vascular responses during 10 min of recording. Data are presented as the individual values and mean ± SEM. * statistically different from 20 Hz stimulation with p<0.05.

First, we ensured the efficiency of the photostimulation paradigm by recording layer II-III pyramidal cells in whole-cell current clamp mode (Fig. 1a). We observed that optogenetic stimulation resulted in the firing of an initial AP that was followed by a train of spikes whose amplitude and frequency transiently decreased before reaching a steady state (Fig. 1a, upper traces). Consistent with the kinetic properties of the H134R ChR2 variant (Lin et al., 2009) and the intrinsic firing properties of pyramidal cells (Karagiannis et al., 2009), the steady-state firing frequency matched the photostimulation frequency up to 5 Hz but was lower at higher frequencies (Fig. 1a, steady-state spike success rate: 100 ± 0 % at 1, 2 and 5 Hz, 70 ± 11 % at 10 Hz and 55 ± 12 % at 20 Hz). These observations demonstrate efficient pyramidal cell activation over a wide range of photostimulation frequencies.

To test the hypothesis that neuronal activity induces vasoconstriction, we analyzed the optogenetically induced response of penetrating arterioles. Layer I arterioles were imaged for 30 minutes in cortical slices (Supplementary Fig.2; Supp. table 1) without preconstriction to facilitate observation of vasoconstriction (Cauli et al., 2004). Examination of the evoked vascular response over 30-minutes (Supplementary Fig.2) showed that increasing the frequency of photostimulation shifted the overall vascular response from a barely discernible delayed response between 1 Hz and 5 Hz to a sustained vasoconstriction at 10 Hz and above which began less than 2 minutes after photostimulation (10 Hz: 1.4 ± 0.4 min; 20 Hz: 1.6 ± 0.5 min). Most vessels (n= 8 of 10 arterioles) showed a strong and rapid vasoconstriction at 20 Hz. On average, this response peaked at 6.8 ± 2.4 min, much earlier than at lower frequencies, which typically required more than 10 min to reach a maximum (Supplementary Fig.2c, 1 Hz: 15.6 ± 4.0 min; 2 Hz: 13.2 ± 2.3 min; 5 Hz: 16.0 ± 3.6 min; 10 Hz: 15.7 ± 2.4 min). Because the vascular response shifted to reliable vasoconstriction, with onset and peak in less than 2 and 10 minutes, respectively, similar to previous observations in cortical slices (Cauli et al., 2004), when the frequency of photostimulation was increased to 20 Hz, we defined the first 10 minutes of recording as the vasoconstriction time frame for subsequent comparisons and analyses. While photostimulation at 1 to 5 Hz failed to elicit fast reliable vascular responses (Fig. 1c-d), 10 Hz photostimulation predominantly induced vasoconstriction (n= 4 of 5 arterioles, Fig. 1 c-d, area under the curve (AUC)= −1.7 ± 1.1 x 10^3^ %.s, n= 5). This response was even more pronounced at 20 Hz, as all arterioles showed vasoconstriction of high magnitude (Fig. 1 b-d; AUC= −3.7 ± 0.7 x 10^3^ %.s, F_(4, 30)_= 6.135, p= 9.89 x 10^-4^, one-way ANOVA, n= 10 arterioles). This difference was particularly striking when comparing the magnitude at 20 Hz (Fig. 1d) with those at 1 Hz (t_(12)_= −3.48, p= 0.0407, t-test), 2 Hz (t_(18)_= −4.09, p= 0.0250, t-test) and 5 Hz (t_(14)_= −3.7, p= 0.0346, t-test). Intense optogenetic stimulation of pyramidal cells has been shown to elicit cortical spreading depression (Chung et al., 2018; Pham et al., 2024), which induces vasoconstriction (Zhang et al., 2024) and fast cell swelling (Zhou et al., 2010). We ruled out this possibility by showing that the rate of change in light transmittance associated with cell swelling remained below that of cortical spreading depression(Zhou et al., 2010) (Supplementary Table 1). On the other hand, ChR2-independent vascular changes induced by high light intensity have been reported (Rungta et al., 2017). We verified that 20 Hz photostimulation did not induce a vascular response in wild-type mice that do not express ChR2 (Supplementary Fig.1). Taken together, our observations indicate that photostimulation of pyramidal cells produces a frequency-dependent vasoconstriction.

### Optogenetic stimulation induces a frequency-dependent, gradual increase in somatic calcium that precedes the vascular response

These observations raise the questions of how pyramidal neurons can induce vasoconstriction at higher AP-firing rates. It is generally accepted that the synthesis and/or release of vasodilatory substances requires an increase in intracellular Ca^2+^ in the releasing cells (Attwell et al., 2010; Cauli and Hamel, 2010), but little is known about the release of vasoconstricting substances. We therefore determined whether an increase in somatic Ca^2+^ concentration in cortical neurons was also dependent on photostimulation frequency. We combined optogenetic stimulation with whole-cell current clamp recording and intracellular Ca^2+^ imaging using Rhod-2 delivered by patch pipette (Fig. 2a). Excitation of this red Ca^2+^ indicator at 585 nm did not induce any voltage response in the recorded pyramidal cells (Fig. 2a), as expected from the action spectrum of ChR2(Lin et al., 2009). In contrast, photostimulation at 470 nm elicited a train of spikes accompanied by a somatic Ca^2+^ increase that decayed for tens of seconds after photostimulation without triggering any significant recurrent spiking activity (Supplementary Fig. 3). The Ca^2+^ response evoked by 20 Hz photostimulation was more than twice of that evoked by 2 Hz photostimulation (Fig. 2b; 2 Hz: ΔF/F_0_= 34.5 ± 3.7 %, n= 9 cells, vs. 20 Hz: ΔF/F_0_= 79.5 ± 17.7 %, n= 9 cells; t _(16)_= 2.485, p= 0.024397), while the average number of evoked spikes was about five times higher (Fig. 1a and supplementary Fig. 3). These results demonstrate a frequency-dependent increase in intracellular Ca^2+^ induced by photostimulation, that precedes vasoconstriction. We therefore aimed to understand the molecular mechanisms linking neuronal activity to vasoconstriction.

**Figure 2:**
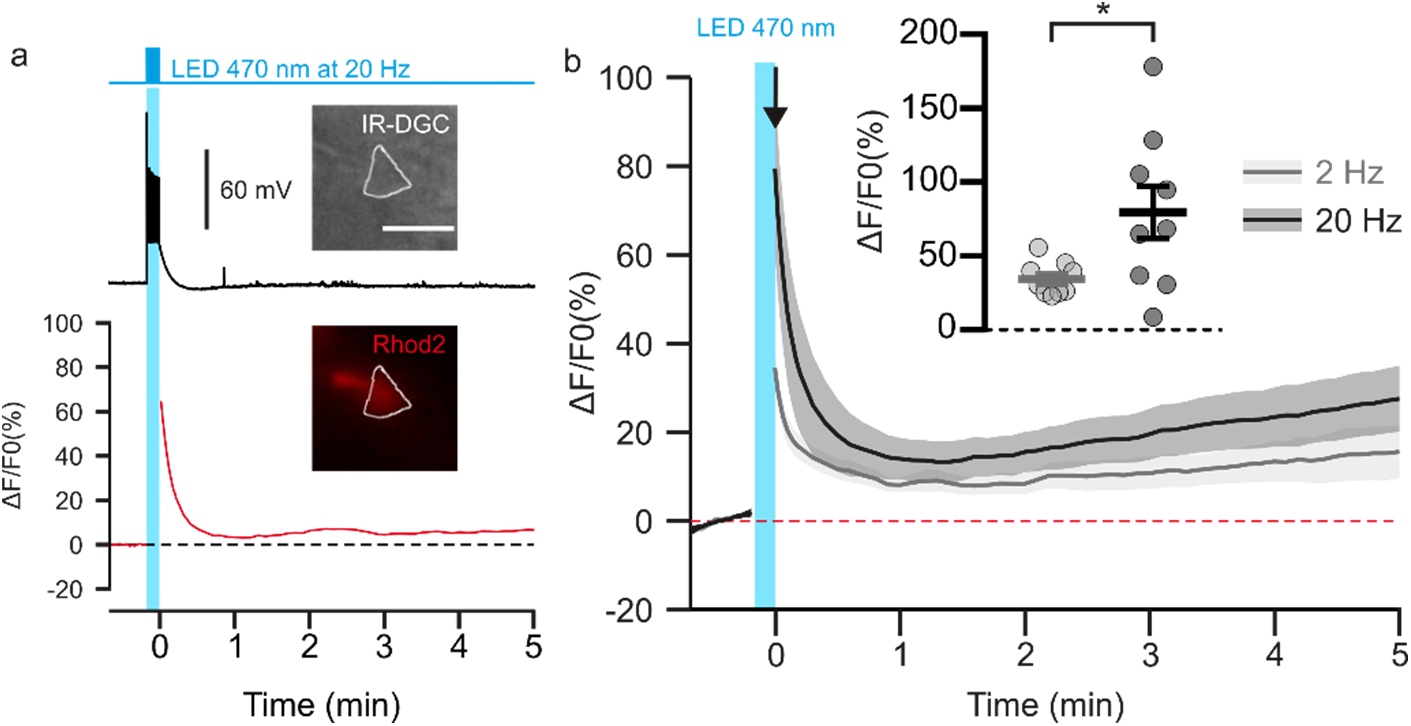
Photostimulation of pyramidal cells elicits a time-locked firing and a frequency-dependent calcium increase. (**a**). Voltage response (top trace) and kinetics of relative fluorescence changes (red bottom trace) induced by photostimulation at 20 Hz. Insets, IR-DGC (top), Rhod2 fluorescence (bottom) pictures of an imaged layer II/III pyramidal cell. The somatic region of interest is outlined in white. Pial surface is upward. Scale bar: 20 µm. **(b)** Mean relative variations of Ca^2+^ fluorescence in response to photostimulation at 2 Hz (grey) and 20 Hz (black). Dashed line represents the baseline. The vertical cyan bar indicates the duration of photostimulation. SEMs envelope the mean traces. Inset, Maximum increase in relative fluorescence changes induced immediately after photostimulation, indicated by the black arrow. The data are shown as the individual values and mean ± SEM. * statistically different with p< 0.05.

### Vasoconstriction induced by pyramidal cells requires AP firing and is partially dependent on glutamatergic transmission

In pyramidal cells, APs induce both somatic Ca^2+^ elevation (Smetters et al., 1999) and glutamate release. To determine whether spiking activity is required for vasoconstriction induced by 20 Hz photostimulation, we blocked APs with the voltage-activated sodium channel blocker tetrodotoxin (TTX, 1 µM, n= 6 arterioles). This treatment completely abolished the vasoconstriction evoked by 20 Hz photostimulation (Fig. 3; AUC= 0.4 ± 0.4 x 10^3^ %.s, t_(14)_= 5.57, p= 8.6656 x 10^-6^). These data indicate that APs are mandatory for neurogenic vascular response and may involve glutamate release.

**Figure 3:**
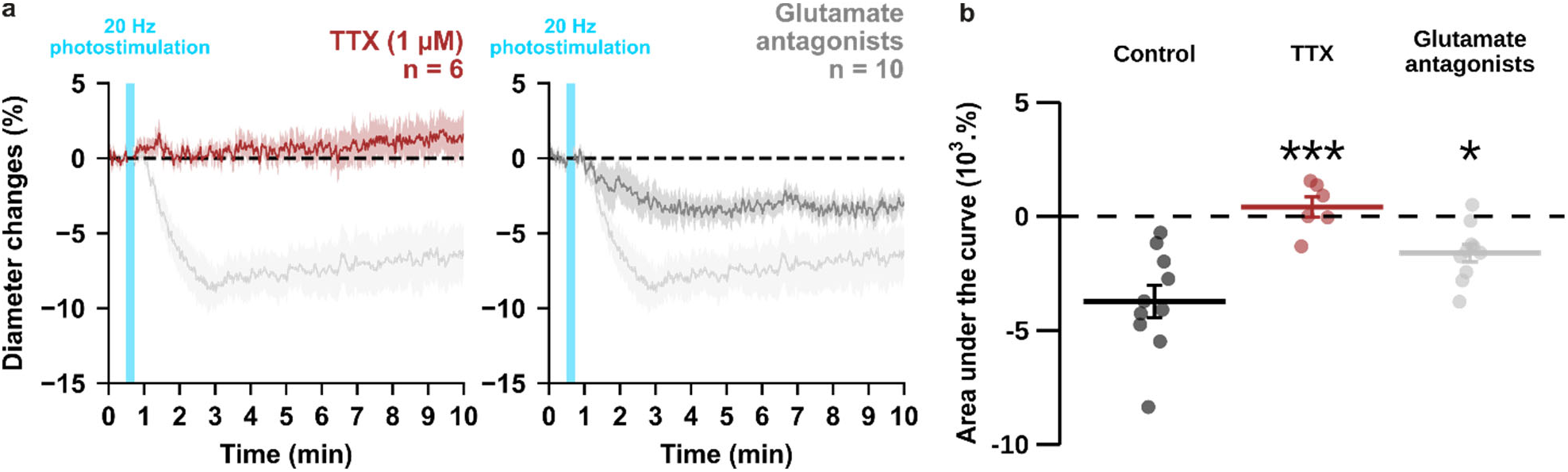
Optogenetically-induced vasoconstriction requires AP firing and partially glutamatergic transmission. Effect of TTX (1 µM, brown, n= 6 arterioles from 5 mice) and cocktail antagonists of AMPA/kainate (DNQX, 10 µM), NMDA (D-AP5, 50 µM), mGluR1 (LY367385, 100 µM) and mGluR5 (MPEP, 50 µM) receptors (gray, n= 10 arterioles from 6 mice) on **(a)** kinetics and **(b)** magnitude of arteriolar vasoconstriction induced by 20 Hz photostimulation (cyan bar). The SEMs envelope the mean traces. Dashed lines represent the initial diameter. The shaded traces correspond to the kinetics of arteriolar vasoconstriction in control condition (Fig. 1c – 20 Hz). Data are presented as the individual values and mean ± SEM. * and *** statistically different from control condition (Fig. 1c – 20 Hz) with p<0.05 and p<0.001, respectively.

Indeed, high levels of glutamate released from pyramidal cells may activate NPY-expressing interneurons or astrocytes through activation of ionotropic or group I metabotropic glutamate receptors (Girouard et al., 2010; Mulligan and MacVicar, 2004; Uhlirova et al., 2016). It may also directly activate NMDA receptors on arteriolar smooth muscle cells, resulting in a large intracellular Ca^2+^ increase and subsequent vasoconstriction (Zhang et al., 2024). To test the hypothesis that glutamate from pyramidal cells, either directly or indirectly, results in vasoconstriction, we blocked glutamatergic transmission by antagonizing AMPA/kainate, NMDA and group I metabotropic receptors expressed by cortical neurons (Tasic et al., 2016; Zeisel et al., 2015) and juvenile astrocytes (Sun et al., 2013). Glutamate receptor antagonists reduced the magnitude of vasoconstriction (−1.6 ± 0.4 x 10^3^ %.s, t_(18)_= 3.28, p= 0.0160) by approximately half (Fig. 3). Taken together, our data suggest that photostimulation of pyramidal cells elicits a frequency-dependent vasoconstriction that requires AP firing and partially involves glutamatergic transmission. We therefore sought to elucidate the glutamate-independent vasoactive pathway underlying this neurogenic vascular response.

### Pyramidal cells express the mRNAs for a cell autonomous PGE2 and PGF2α synthesis

Several arachidonic acid metabolites produced after intracellular Ca^2+^ elevation, including PGF2α, but also PGE2, exert dose-dependent vasoconstrictive effects (Dabertrand et al., 2013; Rosehart et al., 2021; Zonta et al., 2003). These prostaglandins could therefore be progressively released as the frequency of photostimulation and somatic Ca^2+^ increase and thereby promote vasoconstriction. Layer II-III pyramidal cells have been shown to produce PGE2 (Lacroix et al., 2015). To determine whether the synthesizing enzymes of PGE2 and PGF2α are present in pyramidal cells, we performed single-cell RT-PCR after patch-clamp recording (Devienne et al., 2018). Sixteen layer II-III pyramidal cells were visually identified based on the triangular shape of their soma and a prominent apical dendrite. Their glutamatergic phenotype was confirmed both by their stereotypical regular spiking firing pattern (Fig. 4a) and also by the expression of the vesicular glutamate transporter, vGlut1, and neither of the two GABA synthesizing enzymes, thus excluding possible contamination by GABAergic interneurons (Fig. 4b) (Karagiannis et al., 2009). The rate-limiting enzymes of prostaglandin synthesis, cyclooxygenase-1 (COX-1) and −2 (COX-2), were detected in 25% (n= 4 of 16 cells) and 31% (n= 5 of 16 cells) of pyramidal cells, respectively, (Fig. 4b-d) but were never co-expressed (Fig. 4d). Although the differential expression of COX-1 or COX-2 allowed the definition of three non-overlapping molecular subpopulations of pyramidal cells, they did not show distinctive electrophysiological features (supplementary Table 3). The cytosolic enzyme responsible for synthesizing PGE2 (cPGES) was observed in most pyramidal cells (Fig. 4b-c; 88%, n= 14 of 16 cells). In addition, the microsomal PGES, mPGES1 and mPGES2 were detected in 6% (Fig. 4c, n= 1 of 16 cells) and 38% (Fig. 4b,c; n= 6 of 16 cells) of pyramidal cells, respectively, and were always co-expressed with cPGES. The PGF2α terminal-synthesizing enzyme AKR1B3 was observed in the majority of neurons (Fig. 4b-c; n= 11 of 16 cells, 69%). Occasionally, it was co-detected with the prostamide/prostaglandin F synthase (PM-PGFS, Fig. 4c, n= 5 of 16 cells, 31%) and the PGE2 converting enzyme carbonyl reductase 1 (CBR1, Fig. 4c; n= 3 of 16 cells, 19%). CBR1 was consistently detected alongside at least one PGES. Pyramidal cells positive for COX-1 also expressed PGES (Fig. 4d), with most of them also co-expressing PGFS (n=3 out of 4). All neurons positive for COX-2 co-expressed both PGES and PGFS (Fig. 4d). These molecular observations suggest that subpopulations accounting for about half of layer II-III pyramidal cells express all the transcripts necessary for the synthesis of PGE2 and PGF2α.

**Figure 4:**
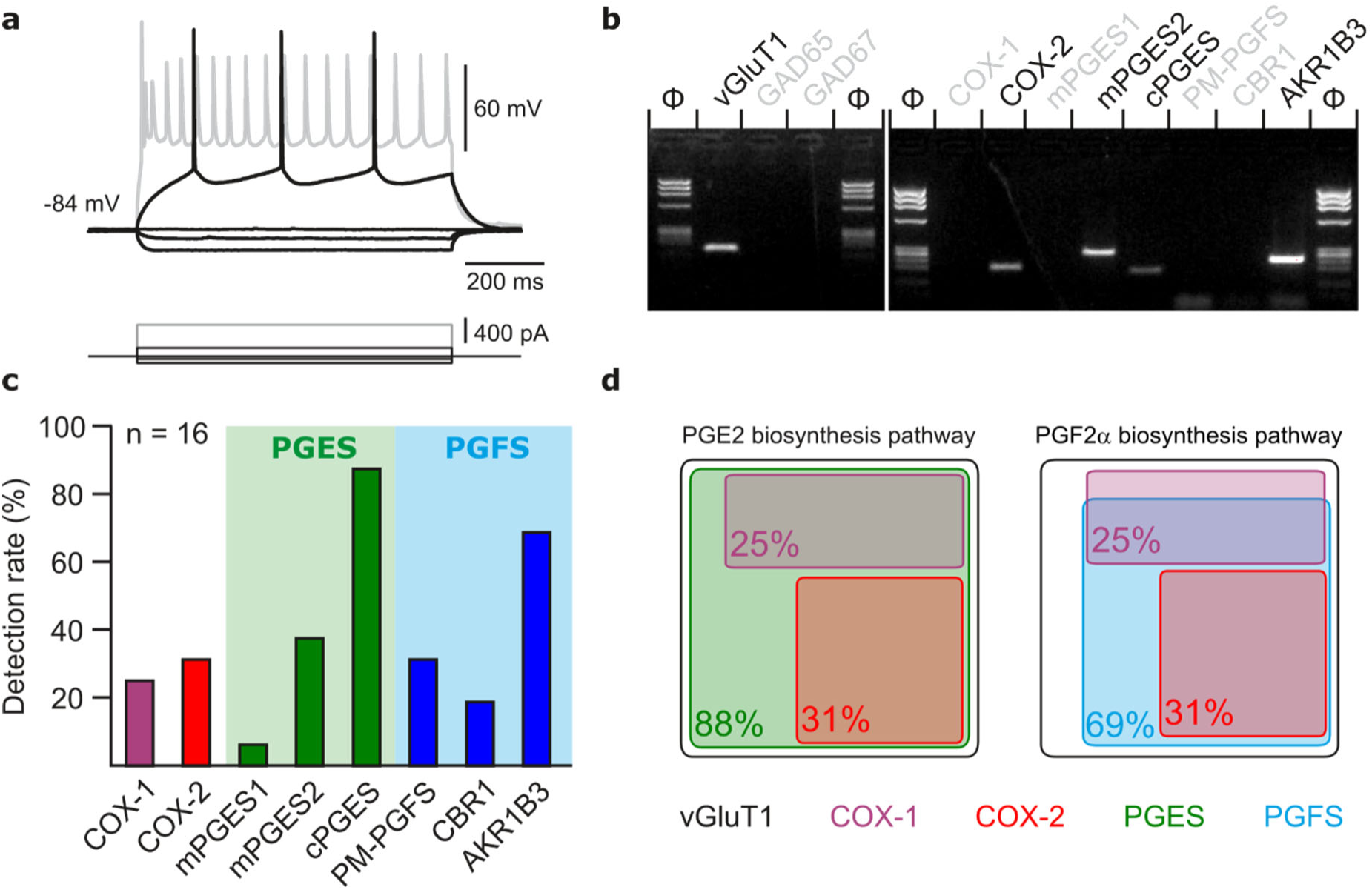
Layer II-III pyramidal cells express PGE2 and PGF2α synthesizing enzymes. **(a)** Voltage responses of a layer II-III pyramidal cell induced by injection of current (bottom traces). In response to a just-above-threshold current pulse, the neuron fired long-lasting action potentials with little frequency adaptation (middle black trace). Near saturation, it exhibits the pronounced spike amplitude accommodation and marked frequency adaptation characteristic of regular spiking cells (upper grey trace). **(b)** Agarose gel analysis of the scRT-PCR products of the pyramidal cell shown in (**a**) revealing expression of vGluT1, COX-2, mPGES2, cPGES, PM-PGFS and CBR1. Φx174 digested by *HaeIII* (Φ) was used as molecular weight marker **(c)** Histogram summarizing the single-cell detection rate of PGE2 and PGf2α synthesizing enzymes in layer II-III pyramidal cells (n= 16 cells from 6 mice). PGES (green zone) corresponds to mPGES1, mPGES2 and/or cPGES and PGFS (blue zone) to PM-PGFS, CBR1 and/or AKR1B3. **(d)** Co-expression of PGE2 and PGf2α synthesizing enzymes in pyramidal cells. The box size is proportional to the detection rate. Note the absence of co-expression between COX-1 (purple) and COX-2 (red). Co-expression of a PGES (left, green) and a PGFS (right, blue) with COX-1 (up) and COX-2 (bottom).

### Prostaglandins underpin vasoconstriction *ex vivo* and *in vivo*

To investigate whether prostaglandins could mediate neurogenic vasoconstriction, we inhibited their synthesis. In cortical slices, the non-selective COX inhibitor indomethacin (5 µM, supplementary Table 4, Fig. 5a and b) completely abolished the vascular response (n= 10 arterioles, AUC= 0.0 ± 0.2 x 10^3^ %.s, t_(18)_= 5.86, p= 3.4385 x 10^-5^). To verify that high-frequency stimulation of pyramidal cells also induces vasoconstriction *in vivo*, 10 Hz photostimulation was reproduced in anesthetized Emx1-cre;Ai32 mice. Pial arterioles diameter measured by 2-photon line scan imaging (Fig. 5c and d) revealed that pyramidal cells induce vasodilation (1^st^ phase) followed by sustained vasoconstriction (2^nd^ phase, Fig. 5d and e). The constriction phase was inhibited by indomethacin (i.v.), indicating the involvement of prostaglandins in the vasoconstriction (AUC_Ctrl_= _-_291.5 ± 92.4 %.s vs. AUC_Indo._= 332.4 ± 184.4 %.s, U_(5,4)_= 0, p= 0.0159, Fig. 5d-f), which confirms our *ex vivo* observations. To determine whether they originated from COX-1 or COX-2 activity, we utilized selective inhibitors in cortical slices. The vasoconstriction magnitude was reduced by the COX-1 inhibitor SC-560 (100 nM, n= 10 arterioles, supplementary Table 4) (−1.4 ± 0.7 x 10^3^ %.s, t_(18)_= 3.54, p= 9.3396 x 10^-3^, Fig. 5a-b). The COX-2 inhibitor NS-398 (10 µM, n= 7 arterioles, supplementary Table 4) completely abolished pyramidal cell-induced vasoconstriction in a more potent manner (Fig. 5a-b; AUC= 0.1 ± 0.3 x 10^3^ %.s, t_(15)_= 5.45, p= 1.0853 x 10^-5^), mimicking the *ex vivo* effect of indomethacin. These observations suggest that prostaglandins, derived mainly from COX-2 activity, and to a lesser extent from COX-1 activity, mediate pyramidal cell-induced vasoconstriction.

**Figure 5:**
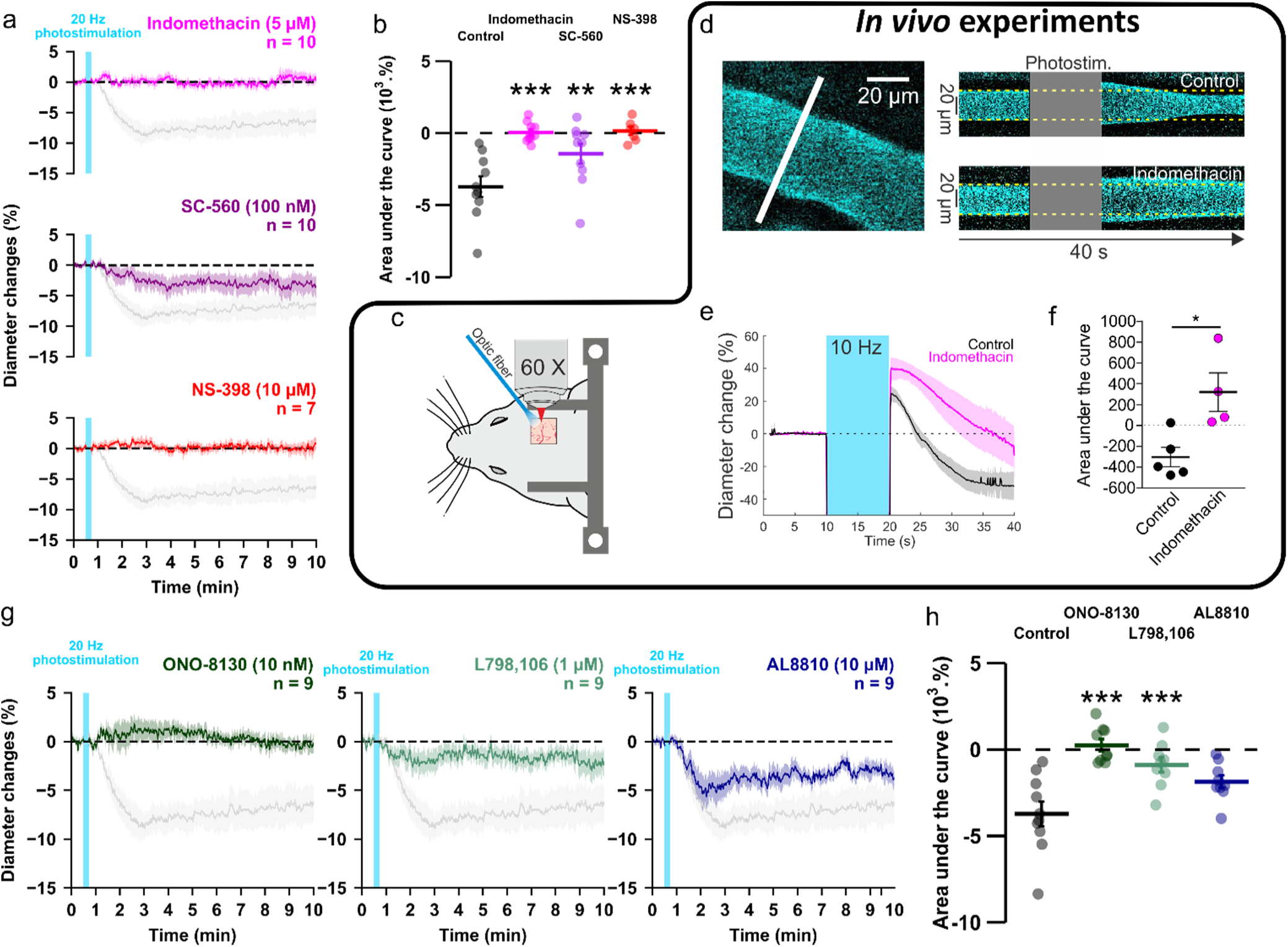
PGE2 mostly derived from COX-2 activity and its EP1 and EP3 receptors mediates vasoconstriction induced by optogenetically activated pyramidal cells. **(a, b)** *Ex vivo* effects of the COX1/2 inhibitor indomethacin (magenta, n= 10 arterioles from 9 mice), the COX-1 inhibitor SC-560 (purple, n= 10 arterioles from 7 mice), and the COX-2 inhibitor NS-398 (red, n= 7 arterioles from 6 mice) on kinetics **(a)** and AUC **(b)** of arteriolar vasoconstriction induced by 20 Hz photostimulation (vertical cyan bar). *In vivo* experiments are highlighted by a black frame. **(c)** Optogenetic stimulation was achieved *in vivo* with an optic fiber through a chronic cranial window over the barrel cortex. **(d)** Left, diameter of pial arterioles labeled with fluorescein dextran (i.v) was measured with line-scan crossing the vessel (white line). Right, Representative examples of vascular response upon photostimulation (10 Hz, 10 s) under control (top) and indomethacin condition (bottom). **(e)** Diameter changes upon photostimulation under control (black; n = 5 arterioles, 4 mice) or indomethacin (magenta; n = 4 arterioles, 4 mice) conditions. **(f)** Area under the curve of the diameter change in control (black) or indomethacin (magenta) conditions calculated between 20 and 40 s (unpaired, two-tailed Mann Whitney test, * p<0.05). **(g, h)** Effects of the EP1, EP3 and FP antagonists, ONO-8130 (10 nM, dark green, n= 9 arterioles from 7 mice), L798,106 (1 µM, light green, n= 9 arterioles from 5 mice) and AL8810 (10 µM, dark blue, n= 9 arterioles from 7 mice), respectively, on kinetics **(g)** and AUC **(h)** of arteriolar vasoconstriction induced by 20 Hz photostimulation. The data are shown as the individual values and mean ± SEM. Dashed line represents the baseline. The SEMs envelope the mean traces. The data are shown as the individual values and mean ± SEM. The shaded traces correspond to the control condition (Fig. 1c – 20 Hz). *, ** and *** statistically different from 20 Hz control condition with p< 0.05, 0.01 and 0.001, respectively.

### PGE2 mediates vasoconstriction by acting primarily on EP1 receptor

To determine the nature of the prostaglandins and their receptors, we selectively antagonized the vasoconstrictor receptors of PGE2, EP1 or EP3, or the FP receptor of PGF2α. The magnitude of vasoconstriction was reduced by the selective EP1 receptor antagonist ONO-8130 (10 nM, n= 9 arterioles, Fig. 5g-h, 0.3 ± 0.3 x 10^3^ %.s, t_(17)_= 6.01, p= 2.8451 x 10^-6^, supplementary Table 4), and to a lesser extent, by the EP3 receptor antagonist L-798,106 (1 µM, n= 9 arterioles, Fig. 5 g-h, −0.9 ± 0.4 x 10^3^ %.s, t_(17)_= 4.30, p= 8.0261 x 10^-4^, supplementary Table 4). Impairing FP receptor signaling with AL-8810 (10 µM, n= 9 arterioles, Fig. 5 g-h, Supplementary table 4) tended to reduce the evoked vasoconstriction, however, it did not reach statistical significance (AUC= −1.9 ± 0.4 x 10^3^ %.s, t_(17)_= 2.82, p= 0.0533). Additionally, the pre-constricted state induced by this weak partial FP agonist(Sharif and Klimko, 2019) (supplementary Fig. 6f) resulted in a diameter reduction of approximately 4% (diameter before application 21.8 ± 2.5 µm vs. during 20.9 ± 2.6 µm, n= 9 arterioles, t_(8)_= 2.374, p= 0.0457, paired t-test), which underestimated the optogenetic vascular response. Taken together, these results indicate that pyramidal cell photoactivation induces vasoconstriction through the release of PGE2 originating mainly from COX-2. This effect primarily acts on the EP1 receptor and, to a lesser extent, on the EP3 receptor.

To test a direct effect of PGE2 through vascular EP1/EP3 activation, we determined whether exogenous agonists of PGE2 receptors could mimic the vasoconstriction induced by pyramidal cell photostimulation. Similar to increasing photostimulation frequencies, exogenous application of PGE2 induced vasoconstriction in a dose-dependent manner which persisted for several minutes after removal (Supplementary Fig. 7a). Likewise, 10 µM sulprostone, an EP1/EP3 agonist with an EC_50_ comparable to that of PGE2 (Boie et al., 1997), mimicked the vasoconstriction induced by 1-10 µM PGE2 (Supplementary Fig. 7b). Application of 10 µM PGE2 in the presence of TTX did not impair the evoked vasoconstriction (Supplementary Fig. 7c). These observations suggest that PGE2 and its EP1 and EP3 receptors mediate a sustained neurogenic vasoconstriction and that once PGE2 is released, its constrictive effect is independent of AP firing.

### Astrocytes through 20-HETE and NPY interneurons are indirect intermediates of pyramidal cell-induced vasoconstriction

In addition to smooth muscle cells, PGE2 released by pyramidal cells can also activate astrocytes and neurons (Clasadonte et al., 2011; Di Cesare et al., 2006), which also express its receptors (Tasic et al., 2016; Zeisel et al., 2015). To assess whether astrocytes could mediate the PGE2-dependent vasoconstriction, we first targeted the large conductance Ca^2+^-activated (BK) channels and the 20-HETE pathways, both of which mediate astrocyte-derived vasoconstriction dependent on glutamatergic transmission (Girouard et al., 2010; Mulligan and MacVicar, 2004). Blockade of BK channels with paxilline (1 µM, n= 10 arterioles, Supplementary table 4) did not impair the vascular response (Fig. 6; AUC = −3.7 ± 0.3 x 10^3^ %.s, t_(18)_ = 0.03, p = 1). Selective inhibition of the 20-HETE synthesizing enzyme, CYP450 ω-hydroxylase, with HET-0016 (100 nM, n= 10 arterioles, Supplementary table 4) reduced the magnitude of the evoked vasoconstriction (Fig. 6; AUC = −1.6 ± 0.7 x 10^3^ %.s, t_(18)_ = 3.32, p = 0.0160). These data suggest that astrocytes partially mediate the vasoconstriction induced by pyramidal cells via 20-HETE but not via K^+^ release. We next determined whether NPY, a potent vasoconstrictor (Cauli et al., 2004), was involved in neurogenic vasoconstriction. Antagonism of the NPY Y1 receptors by BIBP3226 (1 µM, n= 10 arterioles, Supplementary table 4) abolished neurogenic vasoconstriction (Fig. 6; AUC = −0.3 ± 0.2 x 10^3^ %.s, t_(18)_ = 5.28, p = 1.9512 x 10^-5^). These results suggest that neurogenic vasoconstriction induced by pyramidal cell photostimulation involves NPY release and the activation of Y1 receptors (Cauli et al., 2004; Karagiannis et al., 2009; Uhlirova et al., 2016) and astrocytes via 20-HETE in a glutamatergic-dependent and -independent manner.

**Figure 6:**
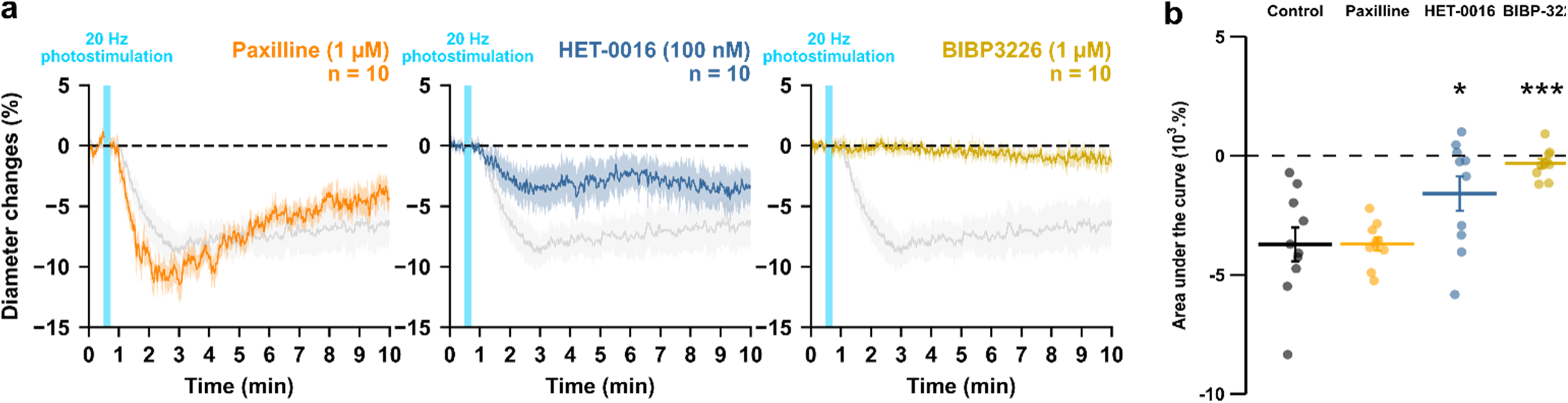
NPY Y1 receptors activation and 20-HETE synthesis mediates the vasoconstriction induced by pyramidal neurons. Effects of paxilline (1 µM, orange, n= 10 arterioles from 6 mice), HET-0016 (100 nM, blue-grey, n= 10 arterioles from 7 mice) and BIBP3226 (1 µM, yellow, n= 10 arterioles from 6 mice) on **(a)** kinetics and **(b)** AUC of arteriolar vasoconstriction induced by 20 Hz photostimulation (vertical blue bar). Dashed line represents the baseline. The SEMs envelope the mean traces. The shaded traces correspond to the control condition (Fig. 1c – 20 Hz). The data are shown as the individual values and mean ± SEM. * and *** statistically different from 20 Hz control condition with p< 0.05 and 0.001.

## Discussion

This study establishes that pyramidal cell activity leads to arteriolar vasoconstriction, and that the magnitude of the vasoconstriction depends on AP firing frequency and correlates with a graded increase in pyramidal cell somatic Ca^2+^. This vascular response partially involves glutamatergic transmission through direct and indirect mechanisms on arteriolar smooth muscle cells. *Ex vivo* and *in vivo* observations revealed that PGE2, predominantly produced by layer II-III COX-2 pyramidal cells, and its EP1 and EP3 receptors play a crucial role in neurogenic vasoconstriction. Pharmacological evidence indicates that some interneurons, via NPY release and activation of Y1 receptors, and to a lesser extent, astrocytes through 20-HETE and possibly COX-1 derived PGE2 play an intermediary role in this process (Figure 7).

**Figure 7:**
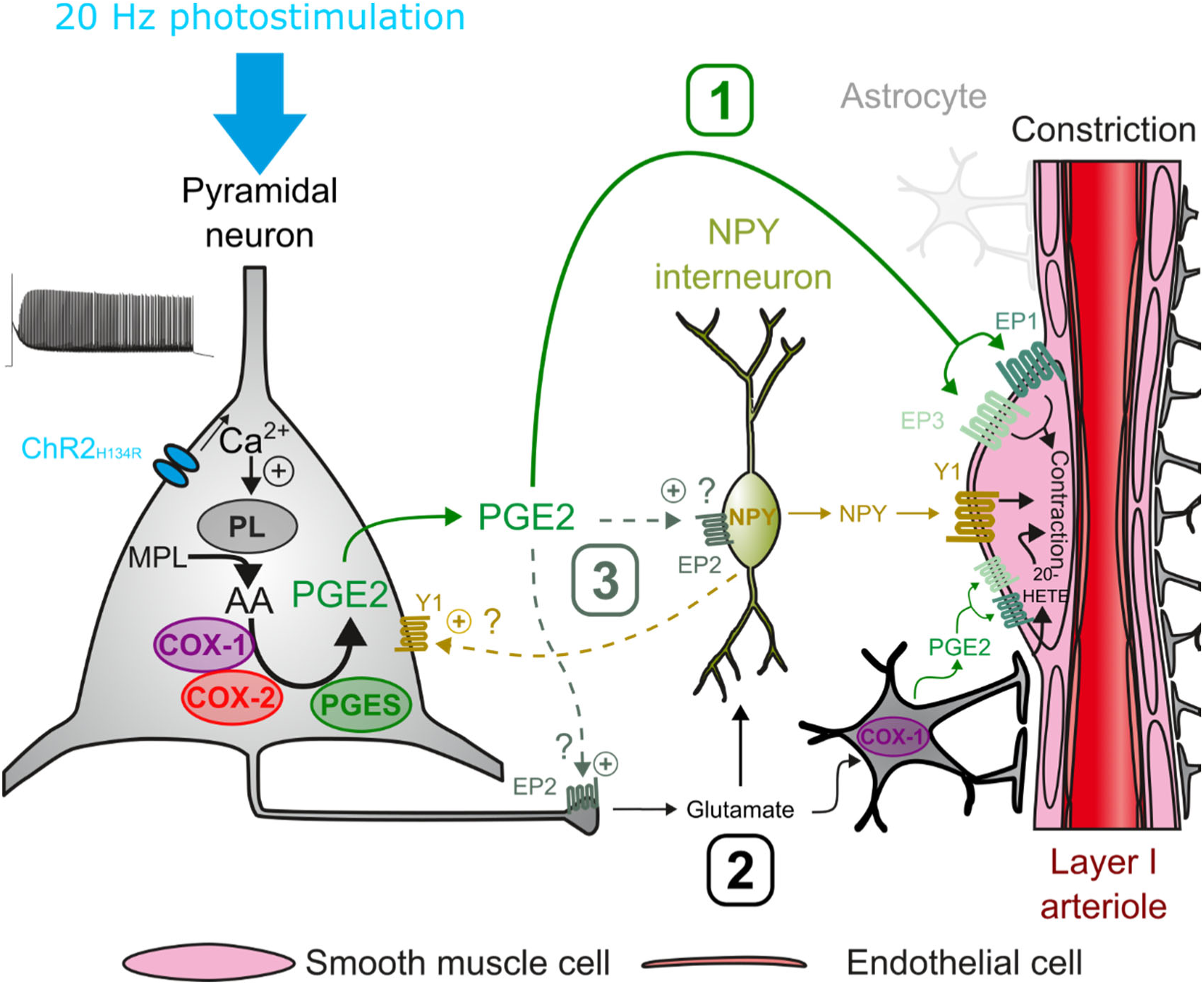
Possible pathways of vasoconstriction induced by pyramidal neurons. 20 Hz photostimulation induces activation of pyramidal neurons expressing channelrhodopsin-2 (ChR2_H134R_) and increases intracellular calcium (Ca^2+^). Arachidonic acid (AA) is released from membrane phospholipids (MPL) by phospholipases (PL) activated by intracellular Ca^2+^ and is metabolized by type-1 and type-2 cyclooxygenases (COX-1 and COX-2) and prostaglandin E2 synthases (PGES) to produce prostaglandin E2 (PGE2). Three non-exclusive pathways can be proposed for arteriolar vasoconstriction in layer I: 1) PGE2 released into the extracellular space may act directly on arteriolar EP1 and EP3 receptors to induce smooth muscle cell constriction. 2) Glutamate released from pyramidal cells may activate neuropeptide Y (NPY) interneurons and NPY is released to act on vascular and neuronal Y1 receptors to constrict smooth muscle cells and promote glutamate release, respectively. Glutamate can also activate astrocytes to induce constriction through the 20-HETE and the COX-1/PGE2 pathways. 3) PGE2 may act on pre- and postsynaptic EP2 receptors to facilitate glutamate release and NPY interneuron activation, respectively.

We found that increasing the frequency of photostimulation in an *ex vivo* preparation caused nearby arteriole to go from a barely discernible response to robust vasoconstriction. In contrast, *in vivo* observations in anesthetized animals, with slower NVC compared to awake animals (Rungta et al., 2021; Uhlirova et al., 2016), have shown that the optogenetic stimulation of pyramidal cells results in a biphasic response: a fast hyperemic/vasodilatory response (Kahn et al., 2013; Lacroix et al., 2015; Scott and Murphy, 2012), which can be followed by a pronounced vasoconstriction (Fig. 5) (Uhlirova et al., 2016). The slow kinetics of the vascular response observed *ex vivo* is comparable with previous observations in slices (Cauli et al., 2004;^,^Rancillac et al., 2006), and is likely due to the lower recording temperature compared to *in vivo*, which slows the synthesis of vasoactive mediators (Rancillac et al., 2006) and downstream reactions. The difficulty in observing vasodilation in cortical slices may be due to relaxed arterioles which favor vasoconstriction (Blanco et al., 2008). The evidence that neurogenic vasoconstriction is frequency-dependent (Fig. 1) and that pharmacologically-induced vasoconstriction persists in preconstructed (Girouard et al., 2010) or pressurized arterioles(Dabertrand et al., 2013), suggests that the neurogenic vasoconstriction primarily depends on a high pyramidal cell firing rate rather than on vascular tone.

Most previous observations did not report a decreased in CBF induced by optogenetic stimulation of pyramidal cells *in vivo* (Lacroix et al., 2015; Scott and Murphy, 2012). This may be attributed to differences in the photostimulation paradigm and/or the specific subtype of pyramidal cells that were stimulated. In our study, we used 10 seconds of photostimulation both *in vivo* and *ex vivo*. Earlier studies have employed shorter photostimulation times, lasting no more than 1 second (Lacroix et al., 2015; Scott and Murphy, 2012; Uhlirova et al., 2016), which may have resulted in an insufficient number of elicited APs to induce robust vasoconstriction. Furthermore, we observed that neurogenic vasoconstriction is highly dependent on COX-2, which is primarily expressed in layer II-III pyramidal cells (Lacroix et al., 2015; Tasic et al., 2016; Zeisel et al., 2015). In our study, photostimulation of almost all pyramidal cells in Emx1-Cre;Ai32 mice (Gorski et al., 2002; Madisen et al., 2012) likely resulted in the release of more COX-2 metabolites. Thy1-ChR2 mice used in previous studies (Scott and Murphy, 2012; Uhlirova et al., 2016), on the other hand, express primarily ChR2 in layer V pyramidal cells (Kahn et al., 2013) which more rarely express COX-2.

Our *ex vivo* and *in vivo* observations revealed that PGE2, primarily derived from COX-2, plays a critical role in neurogenic vasoconstriction by activating EP1 and EP3 receptors expressed by vascular smooth muscle cells (Zhang et al., 2024). Previous studies have shown that COX-2 pyramidal cells, when activated *in vivo* by sensory stimulation or *ex vivo*, induce a NMDA-dependent increase in CBF and vasodilation through PGE2 and EP2/EP4 receptors (Lacroix et al., 2015; Lecrux et al., 2011; Niwa et al., 2000). Differences in the levels and/or sites of action of released PGE2 may explain the absence of secondary vasoconstriction. In Emx1-Cre;Ai32 mice (Gorski et al., 2002; Madisen et al., 2012), optogenetic stimulation may have activated a greater number of COX-2 pyramidal cells and resulted in a higher local release of PGE2 compared to sensory stimulation. Furthermore, since PGE2 is barely catabolized in the cerebral cortex(Alix et al., 2008), most of its removal occurs across the blood-brain barrier by specific transporters. The lack of blood perfusion in brain slices may impair this clearance mechanism, leading to PGE2 accumulation. It is noteworthy that the PGE2-induced vasoconstriction persisted after its removal (supplementary figure 7). A high level of PGE2 may have facilitated the activation of the EP1 receptor, which has a lower affinity than the EP2/EP4 receptors (Boie et al., 1997). Additionally, it may have promoted the rapid desensitization of the dilatory EP4 receptor (Desai et al., 2000) thereby favoring vasoconstriction. Furthermore, PGE2 can induce either EP1-dependent arteriolar dilation or constriction depending on whether it is locally applied to capillaries or arterioles. Constriction prevails when both segments are exposed (Rosehart et al., 2021). Our photostimulation focused on superficial penetrating arterioles, which lack a capillary network in their close vicinity (Kasischke et al., 2011). This may have facilitated the direct EP1-mediated arteriolar constriction (Dabertrand et al., 2013; Rosehart et al., 2021). Overall, these observations suggest that COX-2 pyramidal cells can sequentially promote both vasodilation and vasoconstriction through the release of PGE2, depending on the context.

Consistent with previous reports in rodents (Lacroix et al., 2015; Tasic et al., 2016; Yamagata et al., 1993; Zeisel et al., 2015), the transcripts of the rate-limiting enzymes COX-1 and COX-2, were detected in subpopulations of mouse layer II-III pyramidal cells, respectively. COX-1/2 expression was found to be systematically associated with at least one PGE2 synthesizing enzyme. The major isoforms were cPGES and mPGES2, with the latter being less prevalent (Lacroix et al., 2015; Tasic et al., 2016; Zeisel et al., 2015). The low detection rate of mPGES1, an isoform co-induced with COX-2 by various stimuli(Takemiya et al., 2007; Yamagata et al., 2001), reflects its low constitutive basal expression level. The presence of PM-PGFS, CBR1 and AKR1B3 in layer II-III pyramidal cells is consistent with single-cell RNAseq data (Tasic et al., 2016; Zeisel et al., 2015). The expression of a PGFS was systematically observed in COX-2 positive pyramidal cells and in a majority of COX-1 positive neurons, similar to PGES. These observations collectively indicate that subpopulations of layer II-III pyramidal cells express the mRNAs required for PGE2 and PGF2α synthesis derived from COX-1 or COX-2 activity. Our pharmacological observations did not reveal a contribution of PGF2α in neurogenic vasoconstriction, despite the potential ability of pyramidal cells to produce it. This is likely because PGF2α is only detectable in pyramidal neurons under conditions where COX-2 is over-expressed (Takei et al., 2012).

Pyramidal cells may have an indirect effect on vascular activity through the activation of intermediate cell types, in addition to the direct vascular effects of PGE2 and glutamate (Zhang et al., 2024). Consistent with previous observations, we found that glutamate transmission from pyramidal cells is involved to some extent (Uhlirova et al., 2016). Additionally, we found that the NPY Y1 receptor plays a key role in neurogenic vasoconstriction. It is likely that glutamatergic transmission contributed to NPY release, considering that NPY GABAergic interneurons express a wide range of ionotropic and metabotropic glutamate receptors(Tasic et al., 2016; Zeisel et al., 2015). Consistently, the Y1 receptor has been shown to be involved in vasoconstriction induced by sensory and optogenetic stimulation of GABAergic interneurons (Uhlirova et al., 2016). Activation of group I metabotropic receptors in perivascular astrocytes has been shown to promote vasoconstriction via BK channel-dependent K^+^ release (Girouard et al., 2010) or 20-HETE (Mulligan and MacVicar, 2004). However, the neurogenic vasoconstriction was not affected by the blockade of BK channels, which rules out this astrocytic pathway. In contrast, the inhibition of ω-hydroxylase partially reduced neurogenic vasoconstriction, suggesting the involvement of 20-HETE. Additionally, astrocytes may also have contributed to vasoconstriction through the release of PGE2 derived from COX-1 (Attwell et al., 2016), as indicated by its mild impairment under SC-560.

The observation that both EP1 and Y1 antagonists abolished the vasoconstriction suggests that PGE2 and NPY may act in series and/or in a more complex manner involving their neuronal receptors. One possibility is that PGE2 activates NPY interneurons via the EP1 receptor. However, NPY interneurons barely express its transcript (Tasic et al., 2016; Zeisel et al., 2015) and PGE2 constricts arterioles independently of AP firing, suggesting a direct vascular effect of PGE2. Nevertheless, PGE2 may have facilitated NPY release via pre- and postsynaptic EP2-signaling which have been shown to facilitate glutamate release (Sang et al., 2005) and to induce neuronal firing (Clasadonte et al., 2011), respectively. On the other hand, in addition to smooth muscle cells, Y1 receptors are also enriched in pyramidal (Smith et al., 2019), including COX-2 positive ones (Tasic et al., 2016), and this receptor has been shown to increase extracellular glutamate in the hippocampus (Meurs et al., 2012). By promoting glutamate and possibly PGE2 release, neuronal activation of the Y1 receptor by NPY may also have favored direct (*i.e.* PGE2) and indirect (*i.e.* 20-HETE) vasoconstrictive pathways. The combined activation of vascular and neuronal Y1 receptors may explain the complete blockage of optogenetically induced vasoconstriction by its antagonist BIBP3226. In *ex vivo* relaxed arterioles, where vasoconstriction is favored (Blanco et al., 2008), G_q_ or G_i_ signaling of EP1 or Y1 receptors, respectively, appears sufficient to induce vasoconstriction. *In vivo*, where blood flow both induces myogenic tone and allows PGE2 clearance, NPY and PGE2 could also synergistically promote vasoconstriction by decreasing and increasing cAMP and Ca^2+^ levels, respectively, in smooth muscle cells. PGE2 and NPY may also exert temporally distinct vasoconstrictor effects. Indeed, exogenous application of NPY induces a rapid and transient vasoconstriction that returns to baseline levels after removal (Cauli et al., 2004), whereas PGE2-induced vasoconstriction is slower and more persistent (supplementary figure 7). The more transient effect of NPY likely reflects the presence of multiple NPY-degrading enzymes (Wagner et al., 2015) and/or the desensitization of the Y1 receptor (Gicquiaux et al., 2003, 2002; Tsurumaki et al., 2002) which is not the case for PGE2(Alix et al., 2008) and its vasoconstrictor receptors.

In awake mice, synchronous pyramidal cell activity occurs in the absence of any stimulus during the so-called resting state, but it is observed at a much lower frequency than that which triggers vasoconstriction and is associated with increased blood volume (Ma et al., 2016). Therefore, neurogenic vasoconstriction described here is unlikely to occur under these conditions. Brief sensory stimulation increases pyramidal cell activity and largely causes vasodilation in both awake and anesthetized animals (Rungta et al., 2021). This hyperemic response can be followed by delayed vasoconstriction (Devor et al., 2007) and involves NPY/Y1 receptor signaling (Uhlirova et al., 2016), similar to the mechanisms reported here. It remains unclear whether PGE2 signaling is also involved in this secondary response. During prolonged sensory stimulation the evoked hyperemic response appears to be more restricted to the activated area at the end of the stimulation than at the beginning (Berwick et al., 2008). It is possible that neurogenic vasoconstriction contributes to the later spatial confinement of the vascular response. The time-locked photostimulation of virtually all pyramidal cells leading to vasoconstriction would have resulted in hypersynchrony, a phenomenon that can be observed during sleep/wake transitions (Asadi-Pooya and Sperling, 2019). A decrease in hemodynamics has been reported during the transition from rapid eye movement sleep to wakefulness (Gheres et al., 2023; Tsai et al., 2021), possibly involving neurogenic vasoconstriction. Hypersynchrony is also observed in pathological conditions such as epileptic seizures (Jiruska et al., 2013) and in early stages of Alzheimer’s disease (Bezzina et al., 2015; Palop et al., 2007). Although vasoconstriction observed in epilepsy (Farrell et al., 2016) exhibits similarities to the neurogenic vasoconstriction described herein, there are notable differences between the two. Like neurogenic vasoconstriction, seizure-induced hypoperfusion is dependent on COX-2 (Farrell et al., 2016; Tran et al., 2020) and, to some extent on PGE2 (Farrell et al., 2016), likely through EP1 and/or EP3 receptors. However, epileptic seizures induce the overexpression of both COX-2 and mPGES1 (Takemiya et al., 2007; Yamagata et al., 1993) as well as the ectopic expression of NPY (Baraban, 2004). Similar transcriptional upregulations have also been reported in Alzheimer’s disease (Bezzina et al., 2015; Chaudhry et al., 2008; Palop et al., 2007; Pasinetti and Aisen, 1998) Additionally, PGF2α synthesis by COX-2 pyramidal cells is also observed during seizures (Takei et al., 2012). Taken together, these observations suggest that the mechanisms governing neurogenic vasoconstriction are exacerbated in pathological hypersynchrony and may represent potential therapeutic targets.

This neurogenic vasoconstriction, observed during strong pyramidal cell activity, may seem counterintuitive as it would lead to an undersupply of energy substrates despite a high energy demand. However, vasoconstriction has been reported contralateral to the main activated area, despite bilateral increases in neuronal activity and blood glucose uptake (Devor et al., 2008), suggesting that neurogenic vasoconstriction play a physiological role. Through glutamate uptake by astrocytes, neuronal activity stimulates blood glucose uptake and lactate release (Pellerin and Magistretti, 1994; Voutsinos-Porche et al., 2003). In addition to its role as an oxidative energy substrate for cortical neurons, lactate is also a signaling molecule that enhances their spiking activity (Karagiannis et al., 2021). Therefore, uncontrolled lactate supply and metabolism could potentially lead to deleterious hyperactivity (Cauli et al., 2023; Sada et al., 2015). Thus, the purpose of neurogenic vasoconstriction may be to restrict energy delivery to prevent an overexcitation of the cortical network.

Here, using multidisciplinary approaches, we describe a new mechanism of vasoconstriction that depends on a high firing rate of pyramidal cells. This neurogenic vasoconstriction primarily involves the release of COX-2-derived PGE2 and activation of EP1 and EP3 receptors. It is mediated by direct effects on vascular smooth muscle cells but also by indirect mechanisms involving NPY release from GABAergic interneurons and astrocytes by 20-HETE synthesis. In contrast to previously described mechanisms of neurogenic vasoconstriction, that have been mostly associated with GABAergic interneurons and neuronal inhibition (Cauli et al., 2004; Devor et al., 2007; Krawchuk et al., 2020; Lee et al., 2020; Uhlirova et al., 2016), our data suggest the involvement of glutamatergic excitatory neurons and increased neuronal activity. This finding will help to update the interpretation of the functional brain imaging signals used to map network activity in health and disease (Iadecola, 2017; Zhang and Raichle, 2010). This excitatory form of neurogenic vasoconstriction may also help to understand the etiopathogenesis of epilepsy (Farrell et al., 2016; Tran et al., 2020) and Alzheimer’s disease (Palop and Mucke, 2010) in which increased cortical network activity and hypoperfusion often overlap.

## Materials and methods

### Animals

Homozygous Emx1-Cre mice [Jackson Laboratory, stock #005628, B6.129S2-Emx1*^tm1(cre)Krj^*/J (Gorski et al., 2002)] were crossed with homozygous Ai32 mice [Jackson Laboratory, stock #012569, B6;129S-Gt(ROSA)26Sor^tm32(CAG-COP4*H134R/EYFP)Hze^/J (Madisen et al., 2012)] to obtain heterozygous Emx1^cre/WT^;Ai32^ChR2/WT^ mice for optogenetic stimulations. C57BL/6RJ mice were used for PGE2 and sulprostone exogenous applications, control optogenetic experiments and single-cell RT-PCR. 16-21 postnatal day-old females and males were used for all *ex vivo* experiments. Female Emx1^cre/WT^;Ai32^ChR2/WT^ mice, 3 to 5-month-old, were used for *in vivo* experiments.

All experimental procedures using animals were carried out in strict accordance with French regulations (Code Rural R214/87 to R214/130) and conformed to the ethical guidelines of the European Communities Council Directive of September 22, 2010 (2010/63/UE). Mice were fed *ad libitum* and housed in a 12-hour light/dark cycle. *In vivo* experiments were done in accordance with the Institut national de la santé et de la recherche médicale (Inserm) animal care and approved by the ethical committee Charles Darwin (Comité national de réflexion éthique sur l’expérimentation animale – n°5) (protocol number #27135 2020091012114621).

### *Ex vivo* slice preparation

Mice were deeply anesthetized by isoflurane (IsoVet, Piramal Healthcare UK or IsoFlo, Axience) evaporation in an induction box then euthanized by decapitation. The brain was quickly removed and placed in cold (∼4°C), oxygenated artificial cerebrospinal fluid (aCSF) containing (in mM): 125 NaCl, 2.5 KCl, 1.25 NaH2PO4, 2 CaCl2, 1 MgCl2, 26 NaHCO3, 10 glucose, 15 sucrose and 1 kynurenic acid (Sigma-Aldrich). 300 µm-thick coronal slices containing the barrel cortex were cut with a vibratome (VT1000s; Leica) and were allowed to recover at room temperature for at least 45 min with oxygenated aCSF (95% O2/5% CO2) (Devienne et al., 2018). The slices were then transferred to a submerged recording chamber and perfused continuously at room temperature (20-25 °C) at a rate of 2 ml/min with oxygenated aCSF lacking kynurenic acid.

### Whole-cell recordings

Patch pipettes (5.5 ± 0.2 MΩ) pulled from borosilicate glass were filled with 8 µl of RNase free internal solution containing (in mM): 144 K-gluconate, 3 MgCl2, 0.5 EGTA, 10 HEPES, pH 7.2 (285/295 mOsm). For electrophysiological recordings combined with calcium imaging, EGTA was replaced by 200 µM Rhod-2 (20777, Cayman chemicals). Whole-cell recordings were performed using a patch-clamp amplifier (Axopatch 200B, MDS). Data were filtered at 5-10 kHz and digitized at 50 kHz using an acquisition board (Digidata 1440, MDS) attached to a personal computer running pCLAMP 10.2 software package (MDS). Electrophysiological properties were determined in current-clamp mode (Karagiannis et al., 2009). Membrane potential values were corrected for theoretical liquid junction potential (−15.6 mV). Resting membrane potential of neurons was measured immediately after passing in whole-cell configuration. Only neurons with a resting membrane potential more hyperpolarized than −60 mV were analyzed further.

### Optogenetic stimulation

Optogenetic stimulation was achieved through the objective using a 470 nm light emitting device (LED, CoolLED, Precise Excite) attached to the epifluorescence port of a BX51WI microscope (Olympus) and a set of multiband filters consisting of an excitation filter (HC 392/474/554/635, Semrock), a dichroic mirror (BS 409/493/573/652, Semrock), and an emission filter (HC 432/515/595/730, Semrock). Photostimulation consisted of a 10-s train of 5 ms light pulses at an intensity of 38 mW/mm² and delivered at five different frequencies (1, 2, 5, 10 and 20 Hz).

### Infrared imaging

Blood vessels and cells were observed in slices under infrared illumination with Dodt gradient contrast optics (IR-DGC, Luigs and Neumann) using a double-port upright microscope (BX51WI, Olympus) and a collimated light emitting device (LED; 780 nm; ThorLabs) as the transmitted light source, a 40X (LUMPlanF/IR, 40X/0.80 W, Olympus) or a 60X (LUMPlan FL/IR 60X/0.90 W, Olympus) objective and a digital camera (OrcaFlash 4.0, Hamamatsu) attached to the front port of the microscope. Penetrating arterioles in layer I were selected by IR-DGC videomicroscopy based on their well-defined luminal diameter (10-40 µm), their length remaining in the focal plane for at least 50 µm(Lacroix et al., 2015), and the thickness of their wall (4.1 ± 0.1 µm, n = 176 blood vessels). A resting period of at least 30 min(Zonta et al., 2003) was observed after slice transfer. After light-induced responses, arteriolar contractility was tested by the application of aCSF containing the thromboxane A2 agonist, U46619 (100 nM)(Cauli et al., 2004) or K^+^ enriched solution (composition in mM: 77.5 NaCl, 50 KCl, 1.25 NaH2PO4, 2 CaCl2, 1 MgCl2, 26 NaHCO3, 10 glucose, 15 sucrose). Vessels that did not constrict with these applications were discarded. Only one arteriole was monitored per slice receiving a single optogenetic or pharmacological stimulation. IR-DGC images were acquired at 0.1 Hz for pharmacological applications and at 1 Hz for optogenetic experiments using Imaging Workbench 6.1 software (Indec Biosystems). The focal plane was continuously maintained on-line using IR-DGC images of cells as anatomical landmarks(Lacroix et al., 2015).

### Calcium imaging

Visually and electrophysiologically identified layer II-III pyramidal cells were filled with the calcium-sensitive dye Rhod-2 (200 µM, Cayman chemicals, 20777) using patch pipettes. Optical recordings were made at least 15 min after passing in whole-cell configuration to allow for somatic diffusion of the dye. Rhod-2 was excited with a 585 nm LED (Cool LED, Precise Excite) at an intensity of 0.56 mW/mm² and the filter set used for optogenetic stimulation using the Imaging Workbench 6.1 software (Indec Byosystems). IR-DGC and fluorescence images were acquired by alternating epifluorescence and transmitted light sources. IR-DGC and fluorescence were respectively sampled at 5 Hz and 1 Hz during baseline and optogenetic stimulation, respectively, and at 1 Hz and 0.2 Hz after photostimulation. During photostimulation, bleed-through occurred in the Rhod-2 channel due to the fluorescence of the EYFP-ChR2 transgene (Madisen et al., 2012). Therefore, the Ca^2+^ response could not be reliably analyzed during this period. To compensate for potential x-y drifts, all images were registered off-line using the “StackReg” plug-in (Thévenaz et al., 1998) of the ImageJ 1.53 software. To define somatic regions of interest (ROIs), the soma was manually delineated from IR-DGC images. Fluorescence intensity changes (ΔF/F_0_) were expressed as the ratio (F-F_0_)/F_0_ where F is the mean fluorescence intensity in the ROI at a given time point, and F_0_ is the mean fluorescence intensity in the same ROI during the 30-s control baseline.

### Drugs

All pharmacological compounds were bath applied after a 5-min baseline, and vascular dynamics were recorded during bath application. The following drugs were dissolved in water: D-(-)-2-amino-5-phosphonopentanoic acid (D-AP5, 50 µM, Hellobio, HB0225), 6,7-dinitroquinoxaline-2,3-dione (DNQX, 10µM, Hellobio, HB0262), LY367385 (100µM, Hellobio, HB0398) and BIBP3226 (1 µM, Tocris, 2707). Tetrodotoxin (TTX, 1 µM, L8503, Latoxan) was dissolved in 90 % acetic acid. PGE2 (HB3460, Hellobio), sulprostone (10 µM, Cayman chemical, 14765), 2-methyl-6-(phenylethynyl)pyridine (MPEP, 50 µM, Hellobio, HB0426) and 9,11-dideoxy-9α,11α-methanoepoxy prostaglandin F2α (U-46619, 100 nM, Enzo, BML-PG023) were dissolved in ethanol. Indomethacin (5 µM, Sigma-Aldrich, I7378), SC-560 (100 nM, Sigma-Aldrich, S2064-5MG), NS-398 (10 µM, Enzo, BML-EI261), ONO-8130 (10 nM, Tocris, 5406), L-798,106 (1 µM, Cayman chemical, 11129), AL8810 (10 µM, Cayman chemical, 16735), paxilline (10 µM, Tocris, 2006) and HET0016 (100 µM, Merck, SML2416-5MG) in DMSO. Acetic acid, ethanol and DMSO doses were always used below 0.1%. Synthesis inhibitors and BIBP3226 were applied at least 30 minutes before optogenetic stimulation. TTX was applied at least 15 minutes before, while glutamate receptor antagonists and paxilline were applied at least 10 and 5 minutes before, respectively.

### Vascular reactivity analysis

To compensate for potential x-y drifts, all images were realigned off-line using the “StackReg” plug-in (Thévenaz et al., 1998) of the ImageJ 1.53 software. Luminal diameter was measured in layer I on registered images using custom analysis software developed in MATLAB (MathWorks) (Lacroix et al., 2015). To avoid potential drawbacks due to vessel instability, only arterioles with a stable diameter were analyzed further. Arterioles were considered stable if the relative standard deviation of their diameter during the baseline period was less than 5% (Lacroix et al., 2015). Comparison of the mean arteriolar diameter during the 5-minute baseline and the 5-minute final pharmacological treatments revealed that all drugs, except AL8810, had no effect on resting diameter (Supplementary Fig. 4, 6 and 8).

Diameter changes (ΔD/D_0_) were expressed as (D_t_ – D_0_)/D_0_ where D_t_ is the diameter at the time t and D_0_ is the mean diameter during the baseline period. To eliminate sharp artifacts due to transient loss of focus, diameter change traces were smoothed using a sliding three-point median filter. The overall vascular response over time was captured by the area under the curve of diameter changes after photostimulation. To determine the onset of vasoconstriction, a Z-score was calculated from the diameter change traces using the formula: Z = (x - μ)/ σ, where both the mean μ and the standard deviation σ were calculated from the values before photostimulation. Onset of vasoconstriction was defined as the time after the start of photostimulation at which the Z-score exceeded or fell below −a value of −1.96 (95% criteria) for 10 seconds. If a vessel showed no vasoconstriction, the onset was arbitrarily set at 1800 seconds. Graphs were generated using R software version 4.3.0 (Team et al., 2023) and Matplotlib package (Caswell et al., 2023).

### Intrinsic optical signals analysis

Variations in IR light transmittance (ΔT)(Zhou et al., 2010) were determined using ImageJ 1.53 software according to: ΔT = (T_t_ − T_0_)/ T_0_ where T_t_ is the light transmittance at a time t and T0 is the average light transmittance during the baseline period of a squared region of interest of 100 µm x 100 µm manually delineated in layer I. The rate of ΔT change was determined as the first derivative of ΔT (dΔT/dt, where ΔT is the change in light transmittance and t is time). Slices that showed a maximum rate of increase of dΔT/dt greater than 2%/s, indicating the occurrence of spreading depression (Zhou et al., 2010), were excluded.

### Surgery

Chronic cranial windows were implanted one week after the head bar surgery as previously described (Tournissac et al., 2022). We used a 100 μm thick glass coverslip over the barrel cortex (∼3mm^2^). Before two-photon experiments, a recovery period of 7-10 days minimum was maintained.

### Two-photon imaging and photostimulation

For two-photon excitation, we used a femtosecond laser (Mai Tai eHP; SpectraPhysics) with a dispersion compensation module (Deepsee; SpectraPhysics) emitting 70-fs pulses at 80 MHz. The laser power was attenuated by an acousto-optical modulator (AA Optoelectronic, MT110-B50-A1.5-IR-Hk). Scanning was performed with Galvanometric scanner (GS) mirrors (8315KM60B; Cambridge Technology). Fluorescein was excited at 920 nm and the emitted light was collected with a LUMFLN60XW (Olympus, 1.1 NA) water immersion objective. Collected photons were sorted using a dichroic mirror centered at 570 nm, a FF01-525/25 nm filter (Semrock) and a GaAsP (Hamamatsu) photomultipliers tube. Customized LabView software was used to control the system. Line scans were drawn across pial vessels to measure the change in arterioles diameter, which are not compromised by the fluorescence from the ChR2-EYFP transgene in the parenchyma (Madisen et al., 2012), are less affected by potential movement in the x-y plan than in penetrating arterioles, and whose dilation dynamics are similar in the somatosensory cortex (Rungta et al., 2021).

Mice were anesthetized with a mixture of ketamine and medetomidine (100 and 0.5 mg∕kg, respectively, intraperitoneal (i.p.)) during imaging sessions. Body temperature was maintained at 36.5°C using a retro-controlled heating pad. Fluorescein dextran (70 kDa) was injected i.v. through a retro-orbital injection to label brain vessels. Mice received continuous air through a nose cone supplemented with oxygen to reach a final concentration of 30% O_2_. Photostimulation was delivered with a 473 nm laser (Coblot MLD, Sweden) through an optic fiber placed above the glass coverslip and directed at the pial artery of interest. Each photostimulation consisted of a 10-s train of 5 ms light pulses at an intensity of 1 mW delivered at 10 Hz, with a 5-minute interstimulus interval to allow full recovery to baseline. Indomethacin (10mg/kg, #15425529, Thermo Fisher Scientific) was administered i.v. through a retroorbital injection.

### Imaging analysis

Pial arteriole diameter change was determined with line-scan acquisitions and a home-made Matlab script as previously described (Rungta et al., 2018). Trials from the same vessel were averaged (with a 0.1 s interpolation) for analysis. Area under the curve and statistics were performed using GraphPad Prim (version 6).

### Cytoplasm harvesting and single-cell RT-PCR

At the end of the whole-cell recording, which lasted less than 15 min, the cytoplasmic content was collected in the recording pipette by applying a gentle negative pressure. The pipette’s content was expelled into a test tube and RT was performed in a final volume of 10 µl as described previously (Devienne et al., 2018). The scRT-PCR protocol was designed to probed simultaneously the expression of prostaglandins synthesizing enzymes and neuronal markers (Lacroix et al., 2015). Prostaglandins synthesizing enzymes included COX-1 and COX-2, the terminal PGE2 synthases (PGES): mPGES1, mPGES2 and cPGES, the terminal PGF2α synthases (PGFS): PM-PGFS (Prxl2b) and AKR1B3 and the carbonyl reductase CBR1. Neuronal markers included the vesicular glutamate transporter, vGluT1, and the two isoforms of glutamic acid decarboxylase, GAD65 and GAD67. Two-step amplification was performed essentially as described (Devienne et al., 2018). First, cDNAs present in the 10 µl reverse transcription reaction were simultaneously amplified with all external primer pairs listed in Supplementary table 2. Taq polymerase (2.5 U; Qiagen) and external primers mix (20 pmol each) were added to the manufacturer’s buffer (final volume, 100 µl), and 20 cycles (95◦C, 30 s; 60◦C, 30 s; and 72◦C, 35 s) of PCR were performed. Second rounds of PCR were performed using 1 µl of the first PCR product as a template. In this second round, each cDNA was amplified individually using its specific nested primer pair (Supplementary table 2) by performing 35 PCR cycles (as described above). 10 µl of each individual PCR product were run on a 2% agarose gel stained with ethidium bromide using ΦX174 digested by *HaeIII* as a molecular weight marker. The efficiency of the protocol was validated using 500 pg of total forebrain RNAs (Supplementary Fig. 5).

### Statistical analyses

Statistical analyses were performed using GraphPad Prism version 7.00 for Windows (GraphPad Software, La Jolla California USA, www.graphpad.com) and R software version 4.3.0 (Team et al., 2023). Normality of distribution was assessed using the Shapiro-Wilk tests. Equality of variance was assessed using Brown-Forsythe tests for comparisons between groups and using F-tests for comparisons with a control group. Parametric tests were only used if these criteria were met. Statistical significance of morphological and physiological properties of penetrating arterioles was determined using one-way ANOVA for comparison between groups. Statistical significance of calcium was determined using two-tailed unpaired t-tests and Statistical significance of vascular responses were appreciated using Tukey posthoc tests for the different frequencies conditions and using Dunnett’s posthoc tests for the different pharmacological conditions compared to the 20 Hz condition without pharmacological compound. False discovery rate correction was used for multiple comparisons. Statistical significance of vascular diameter for drug applications was determined using two-tailed paired t-tests. Statistical significance on all figures uses the following convention: *p<0.05, **p<0.01 and ***p<0.001.

## Data availability statement

The data that support the results of this study are available from the corresponding author upon reasonable request.

## Acknowledgements

The authors thank Dr Rebecca Piskorowski for constructive criticism of the manuscript. We acknowledge the invaluable support of the animal facilities of IBPS (RongIBPS) and Institut de la vision for their expert care and maintenance of the animals used in this study. Financial support was provided by grants from the Agence Nationale pour la Recherche (ANR-17-CE37-0010-03, B.C.; CE37_2020_TF-fUS-CADASIL, S.C.; ANR-20-CE14-0025, D.L.; ANR-23-CE14-0038-01, B.C.), the Fondation Alzheimer France (M21JRCN009, S.C.) and the i-Bio initiative of Sorbonne University (B.C.). B.L.G. and E.B. were supported by fellowships from Fondation pour la Recherche sur Alzheimer and M.T. by a fellowship from the Fondation pour la Recherche Médicale (SPF201909009103).

## Author contributions

B.L.G., M.T., and E.B. designed experiments, acquired, and analyzed the data, and edited the manuscript. S.P. acquired the data. B.L.G. and B.C. drafted the manuscript. I.D. edited the manuscript. H.S. analyzed the data. D.L. designed experiments and edited the manuscript. S.C. and B.C. designed experiments and edited the manuscript.

## References

Alix E, Schmitt C, Strazielle N, Ghersi-Egea JF. 2008. Prostaglandin E2 metabolism in rat brain: Role of the blood-brain interfaces. Cerebrospinal Fluid Res 5. doi:10.1186/1743-8454-5-5

Asadi-Pooya AA, Sperling MR. 2019. Normal Awake, Drowsy, and Sleep EEG Patterns That Might Be Overinterpreted as Abnormal. Journal of Clinical Neurophysiology. doi:10.1097/WNP.0000000000000585

Attwell D, Buchan AM, Charpak S, Lauritzen M, Macvicar BA, Newman EA. 2010. Glial and neuronal control of brain blood flow. Nature 468:232–243. doi:10.1038/nature09613

Attwell D, Mishra A, Hall CN, O’Farrell FM, Dalkara T. 2016. What is a pericyte? Journal of Cerebral Blood Flow and Metabolism 36:451–455. doi:10.1177/0271678X15610340

Baraban SC. 2004. Neuropeptide Y and epilepsy: Recent progress, prospects and controversies. Neuropeptides. doi:10.1016/j.npep.2004.04.006

Berwick J, Johnston D, Jones M, Martindale J, Martin C, Kennerley AJ, Redgrave P, Mayhew JEW. 2008. Fine detail of neurovascular coupling revealed by spatiotemporal analysis of the hemodynamic response to single whisker stimulation in rat barrel cortex. J Neurophysiol 99:787–798. doi:10.1152/jn.00658.2007

Bezzina C, Verret L, Juan C, Remaud J, Halley H, Rampon C, Dahan L. 2015. Early onset of hypersynchronous network activity and expression of a marker of chronic seizures in the Tg2576 mouse model of Alzheimer’s disease. PLoS One 10. doi:10.1371/journal.pone.0119910

Blanco M, Stern JE, Filosa JA. 2008. Tone-dependent vascular responses to astrocyte-derived signals. American journal of physiology 2855–2863. doi:10.1152/ajpheart.91451.2007.

Boie Y, Stocco R, Sawyer N, Slipetz DM, Ungrin MD, Neuschafer-rube F, Puschel GP. 1997. Molecular cloning and characterization of the four rat prostaglandin E 2 prostanoid receptor subtypes 227–241.

Caswell TA, Andrade ES de, Lee A, Droettboom M, Hoffmann T, Klymak J, Hunter J, Firing E, Stansby D, Varoquaux N, Nielsen JH, Gustafsson O, Root B, May R, Elson P, Seppänen JK, Lee J-J, Dale D, Sunden K, hannah, McDougall D, Straw A, Hobson P, Lucas G, Gohlke C, Vincent AF, Yu TS, Ma E, Silvester S, Moad C. 2023. matplotlib/matplotlib: REL: v3.7.2. doi:10.5281/ZENODO.8118151

Cauli B, Dusart I, Li D. 2023. Lactate as a determinant of neuronal excitability, neuroenergetics and beyond. Neurobiol Dis 184:106207. doi:10.1016/j.nbd.2023.106207

Cauli B, Hamel E. 2010. Revisiting the role of neurons in neurovascular coupling. Front Neuroenergetics 2:9. doi:10.3389/fnene.2010.00009

Cauli B, Tong XK, Rancillac A, Serluca N, Lambolez B, Rossier J, Hamel E. 2004. Cortical GABA interneurons in neurovascular coupling: Relays for subcortical vasoactive pathways. Journal of Neuroscience 24:8940–8949. doi:10.1523/JNEUROSCI.3065-04.2004

Chaudhry UA, Zhuang H, Crain BJ, Doré S. 2008. Elevated microsomal prostaglandin-E synthase-1 in Alzheimer’s disease. Alzheimer’s and Dementia 4:6–13. doi:10.1016/j.jalz.2007.10.015

Chung DY, Sadeghian H, Qin T, Lule S, Lee H, Karakaya F, Goins S, Oka F, Yaseen MA, Houben T, Tolner EA, Van Den Maagdenberg AMJMJMJM, Whalen MJ, Sakadžić S, Ayata C. 2018. Determinants of Optogenetic Cortical Spreading Depolarizations. Cerebral Cortex 29:1–12. doi:10.1093/cercor/bhy021

Clasadonte J, Poulain P, Hanchate NK, Corfas G, Ojeda SR, Prevot V. 2011. Prostaglandin E 2 release from astrocytes triggers gonadotropin-releasing hormone (GnRH) neuron firing via EP2 receptor activation. Proc Natl Acad Sci U S A 108:16104–16109. doi:10.1073/pnas.1107533108

Dabertrand F, Hannah RM, Pearson JM, Hill-Eubanks DC, Brayden JE, Nelson MT. 2013. Prostaglandin E2, a postulated astrocyte-derived neurovascular coupling agent, constricts rather than dilates parenchymal arterioles. Journal of Cerebral Blood Flow and Metabolism 33:479–482. doi:10.1038/jcbfm.2013.9

Desai S, April H, Nwaneshiudu C, Ashby B. 2000. Comparison of agonist-induced internalization of the human EP2 and EP4 prostaglandin receptors: Role of the carboxyl terminus in EP4 receptor sequestration. Mol Pharmacol 58:1279–1286. doi:10.1124/mol.58.6.1279

Devienne G, Le Gac B, Piquet J, Cauli B. 2018. Single Cell Multiplex Reverse Transcription Polymerase Chain Reaction After Patch-Clamp. Journal of Visualized Experiments 136:1–12. doi:10.3791/57627

Devor A, Hillman EMC, Tian P, Waeber C, Teng IC, Ruvinskaya L, Shalinsky MH, Zhu H, Haslinger RH, Narayanan SN, Ulbert I, Dunn AK, Lo EH, Rosen BR, Dale AM, Kleinfeld D, Boas DA. 2008. Stimulus-Induced Changes in Blood Flow and 2-Deoxyglucose Uptake Dissociate in Ipsilateral Somatosensory Cortex. Journal of Neuroscience 28:14347–14357. doi:10.1523/JNEUROSCI.4307-08.2008

Devor A, Tian P, Nishimura N, Teng IC, Hillman EMC, Narayanan SN, Ulbert I, Boas DA, Kleinfeld D, Dale AM. 2007. Suppressed Neuronal Activity and Concurrent Arteriolar Vasoconstriction May Explain Negative Blood Oxygenation Level-Dependent Signal. Journal of Neuroscience 27:4452–4459. doi:10.1523/JNEUROSCI.0134-07.2007

Di Cesare A, Del Piccolo P, Zacchetti D, Grohovaz F. 2006. EP2 receptor stimulation promotes calcium responses in astrocytes via activation of the adenylyl cyclase pathway. Cellular and Molecular Life Sciences 63:2546–2553. doi:10.1007/s00018-006-6262-9

Farrell JS, Gaxiola-Valdez I, Wolff MD, David LS, Dika HI, Geeraert BL, Wang R, Singh S, Spanswick SC, Dunn JF, Antle MC, Federico P, Campbell Teskey G. 2016. Postictal behavioural impairments are due to a severe prolonged hypoperfusion/ hypoxia event that is COX-2 dependent. doi:10.7554/eLife.19352.001

Gheres KW, Ünsal HS, Han X, Zhang Q, Turner KL, Zhang N, Drew PJ. 2023. Arousal state transitions occlude sensory-evoked neurovascular coupling in neonatal mice. Commun Biol 6. doi:10.1038/s42003-023-05121-5

Gicquiaux H, Lecat S, Gaire M, Dieterlen A, Mély Y, Takeda K, Bucher B, Galzi JL. 2003. Neuropeptide Y-induced contraction and its desensitization through the neuropeptide Y receptor subtype in several rat veins. J Cardiovasc Pharmacol 41 **Suppl 1**:S23–7.

Gicquiaux H, Lecat S, Gaire M, Dieterlen A, Mély Y, Takeda K, Bucher B, Galzi JL. 2002. Rapid internalization and recycling of the human neuropeptide Y Y1 receptor. Journal of Biological Chemistry 277:6645–6655. doi:10.1074/jbc.M107224200

Girouard H, Bonev AD, Hannah RM, Meredith A, Aldrich RW, Nelson MT. 2010. Astrocytic endfoot Ca2+ and BK channels determine both arteriolar dilation and constriction. Proc Natl Acad Sci U S A 107:3811–3816. doi:10.1073/pnas.0914722107

Gordon GRJ, Choi HB, Rungta RL, Ellis-Davies GCR, MacVicar BA. 2008. Brain metabolism dictates the polarity of astrocyte control over arterioles. Nature 456:745–9. doi:10.1038/nature07525

Gorski JA, Talley T, Qiu M, Puelles L, Rubenstein JLRR, Jones KR. 2002. Cortical excitatory neurons and glia, but not GABAergic neurons, are produced in the Emx1-expressing lineage. Journal of Neuroscience 22:6309–6314. doi:10.1523/jneurosci.22-15-06309.2002

Grutzendler J, Nedergaard M. 2019. Cellular Control of Brain Capillary Blood Flow: In Vivo Imaging Veritas. Trends Neurosci. doi:10.1016/j.tins.2019.05.009

Hartmann DA, Berthiaume AA, Grant RI, Harrill SA, Koski T, Tieu T, McDowell KP, Faino A V., Kelly AL, Shih AY. 2021. Brain capillary pericytes exert a substantial but slow influence on blood flow. Nat Neurosci 24:633–645. doi:10.1038/s41593-020-00793-2

Hill RA, Tong L, Yuan P, Murikinati S, Gupta S, Grutzendler J. 2015. Regional Blood Flow in the Normal and Ischemic Brain Is Controlled by Arteriolar Smooth Muscle Cell Contractility and Not by Capillary Pericytes. Neuron 87:95–110. doi:10.1016/j.neuron.2015.06.001

Iadecola C. 2017. The Neurovascular Unit Coming of Age: A Journey through Neurovascular Coupling in Health and Disease. Neuron 96:17–42. doi:10.1016/j.neuron.2017.07.030

Iadecola C, Nedergaard M. 2007. Glial regulation of the cerebral microvasculature. Nat Neurosci 10:1369–1376. doi:10.1038/nn2003

Jiruska P, de Curtis M, Jefferys JGR, Schevon CA, Schiff SJ, Schindler K. 2013. Synchronization and desynchronization in epilepsy: Controversies and hypotheses. Journal of Physiology. doi:10.1113/jphysiol.2012.239590

Kahn I, Knoblich U, Desai M, Bernstein J, Graybiel AM, Boyden ES, Buckner RL, Moore CI. 2013. Optogenetic drive of neocortical pyramidal neurons generates fMRI signals that are correlated with spiking activity. Brain Res 1511:33–45. doi:10.1016/j.brainres.2013.03.011

Karagiannis A, Gallopin T, Dávid C, Battaglia D, Geoffroy H, Rossier J, Hillman EMC, Staiger JF, Cauli B. 2009. Classification of NPY-expressing neocortical interneurons. J Neurosci 29:3642–59. doi:10.1523/JNEUROSCI.0058-09.2009

Karagiannis A, Gallopin T, Lacroix A, Plaisier F, Piquet J, Geoffroy H, Hepp R, Naudé J, Le Gac B, Egger R, Lambolez B, Li D, Rossier J, Staiger JF, Imamura H, Seino S, Roeper J, Cauli B. 2021. Lactate is an energy substrate for rodent cortical neurons and enhances their firing activity. Elife 10:1–40. doi:10.7554/elife.71424

Kasischke KA, Lambert EM, Panepento B, Sun A, Gelbard HA, Burgess RW, Foster TH, Nedergaard M. 2011. Two-photon NADH imaging exposes boundaries of oxygen diffusion in cortical vascular supply regions. Journal of Cerebral Blood Flow and Metabolism 31:68–81. doi:10.1038/jcbfm.2010.158

Krawchuk MB, Ruff CF, Yang X, Ross SE, Vazquez AL. 2020. Optogenetic assessment of VIP, PV, SOM and NOS inhibitory neuron activity and cerebral blood flow regulation in mouse somato-sensory cortex. Journal of Cerebral Blood Flow and Metabolism 40:1427–1440. doi:10.1177/0271678X19870105

Lacroix A, Toussay X, Anenberg E, Lecrux C, Ferreirós N, Karagiannis A, Plaisier F, Chausson P, Jarlier F, Burgess SA, Hillman EMCCC, Tegeder I, Murphy TH, Hamel E, Cauli B, Ferreiro N, Burgess SA, Hillman EMCCC, Tegeder I, Murphy TH, Hamel E, Cauli B. 2015. COX-2-Derived Prostaglandin E2 Produced by Pyramidal Neurons Contributes to Neurovascular Coupling in the Rodent Cerebral Cortex. J Neurosci 35:11791–11810. doi:10.1523/JNEUROSCI.0651-15.2015

Lecrux C, Toussay X, Kocharyan A, Fernandes P, Neupane S, Levesque M, Plaisier F, Shmuel A, Cauli B, Hamel E. 2011. Pyramidal Neurons Are “Neurogenic Hubs” in the Neurovascular Coupling Response to Whisker Stimulation. Journal of Neuroscience 31:9836–9847. doi:10.1523/JNEUROSCI.4943-10.2011

Lee JMJ, Stile CL, Bice AR, Rosenthal ZP, Yan P, Snyder AZ, Lee JMJ, Bauer AQ. 2020. Opposed hemodynamic responses following increased excitation and parvalbumin-based inhibition. Journal of Cerebral Blood Flow and Metabolism. doi:10.1177/0271678X20930831

Lin JY, Lin MZ, Steinbach P, Tsien RY. 2009. Characterization of engineered channelrhodopsin variants with improved properties and kinetics. Biophys J 96:1803–1814. doi:10.1016/j.bpj.2008.11.034

Ma Y, Shaik MA, Kozberg MG, Kim SH, Portes JP, Timerman D. 2016. Resting-state hemodynamics are spatiotemporally coupled to synchronized and symmetric neural activity in excitatory neurons. Proceedings of the National Academy of Sciences. doi:10.1073/pnas.1525369113

Madisen L, Mao T, Koch H, Zhuo J, Berenyi A, Fujisawa S, Hsu Y-W a, Garcia AJ, Gu X, Zanella S, Kidney J, Gu H, Mao Y, Hooks BM, Boyden ES, Buzsáki G, Ramirez JM, Jones AR, Svoboda K, Han X, Turner EE, Zeng H. 2012. A toolbox of Cre-dependent optogenetic transgenic mice for light-induced activation and silencing. Nat Neurosci 15:793–802. doi:10.1038/nn.3078

Meurs A, Portelli J, Clinckers R, Balasubramaniam A, Michotte Y, Smolders I. 2012. Neuropeptide Y increases in vivo hippocampal extracellular glutamate levels through Y1 receptor activation. Neurosci Lett 510:143–147. doi:10.1016/j.neulet.2012.01.023

Mishra A, Reynolds JP, Chen Y, Gourine A V., Rusakov DA, Attwell D. 2016. Astrocytes mediate neurovascular signaling to capillary pericytes but not to arterioles. Nat Neurosci 19:1619– 1627. doi:10.1038/nn.4428

Mulligan SJ, MacVicar BA. 2004. Calcium transients in astrocyte endfeet cause cerebrovascular constrictions. Nature 431:195–199. doi:10.1038/nature02827

Niwa K, Araki E, Morham SG, Ross ME, Iadecola C. 2000. Cyclooxygenase-2 contributes to functional hyperemia in whisker-barrel cortex. Journal of Neuroscience 20:763–770. doi:10.1523/jneurosci.20-02-00763.2000

O’Herron P, Chhatbar PY, Levy M, Shen Z, Schramm AE, Lu Z, Kara P. 2016. Neural correlates of single-vessel haemodynamic responses in vivo. Nature 534:378–382. doi:10.1038/nature17965

Palop JJ, Chin J, Roberson ED, Wang J, Thwin MT, Bien-Ly N, Yoo J, Ho KO, Yu GQ, Kreitzer A, Finkbeiner S, Noebels JL, Mucke L. 2007. Aberrant Excitatory Neuronal Activity and Compensatory Remodeling of Inhibitory Hippocampal Circuits in Mouse Models of Alzheimer’s Disease. Neuron 55:697–711. doi:10.1016/j.neuron.2007.07.025

Palop JJ, Mucke L. 2010. Amyloid-Β-induced neuronal dysfunction in Alzheimer’s disease: From synapses toward neural networks. Nat Neurosci 13:812–818. doi:10.1038/nn.2583

Pasinetti GM, Aisen PS. 1998. Cyclooxygenase-2 expression is increased in frontal cortex of Alzheimer’s disease brain. Neuroscience 87:319–324. doi:10.1016/S0306-4522(98)00218-8

Pellerin L, Magistretti PJ. 1994. Glutamate uptake into astrocytes stimulates aerobic glycolysis: A mechanism coupling neuronal activity to glucose utilization. Proc Natl Acad Sci U S A 91:10625–10629. doi:10.1073/pnas.91.22.10625

Pham C, Komaki Y, Deàs-Just A, Le Gac B, Mouffle C, Franco C, Chaperon A, Vialou V, Tsurugizawa T, Cauli B, Li D. 2024. Astrocyte aquaporin mediates a tonic water efflux maintaining brain homeostasis. Elife 13. doi:10.7554/eLife.95873

Rancillac A, Rossier J, Guille M, Tong X-K, Geoffroy H, Amatore C, Arbault S, Hamel E, Cauli B. 2006. Glutamatergic Control of Microvascular Tone by Distinct GABA Neurons in the Cerebellum. J Neurosci 26:6997–7006. doi:10.1523/JNEUROSCI.5515-05.2006

Rosehart AC, Longden TA, Weir N, Fontaine JT, Joutel A, Dabertrand F. 2021. Prostaglandin E2 Dilates Intracerebral Arterioles When Applied to Capillaries: Implications for Small Vessel Diseases. Front Aging Neurosci 13:1–11. doi:10.3389/fnagi.2021.695965

Rungta RL, Chaigneau E, Osmanski BF, Charpak S. 2018. Vascular Compartmentalization of Functional Hyperemia from the Synapse to the Pia. Neuron 1–14. doi:10.1016/j.neuron.2018.06.012

Rungta RL, Osmanski B-F, Boido D, Tanter M, Charpak S. 2017. Light controls cerebral blood flow in naive animals. Nat Commun 8:14191. doi:10.1038/ncomms14191

Rungta RL, Zuend M, Aydin A, Weber B, Charpak S, Boido D. 2021. vascular arbors in layer II / III somatosensory cortex. Commun Biol. doi:10.1038/s42003-021-02382-w

Sada N, Lee S, Katsu T, Otsuki T, Inoue T. 2015. Targeting LDH enzymes with a stiripentol analog to treat epilepsy. Science (1979) 347:1362–1367. doi:10.1126/science.aaa1299

Sang N, Zhang J, Marcheselli V, Bazan NG, Chen C. 2005. Postsynaptically Synthesized Prostaglandin E2 (PGE2) Modulates Hippocampal Synaptic Transmission via a Presynaptic PGE2 EP2 Receptor. Journal of Neuroscience 25:9858–9870. doi:10.1523/JNEUROSCI.2392-05.2005

Schmid F, Barrett MJPP, Jenny P, Weber B. 2019. Vascular density and distribution in neocortex. Neuroimage 197:792–805. doi:10.1016/j.neuroimage.2017.06.046

Scott NA, Murphy TH. 2012. Hemodynamic responses evoked by neuronal stimulation via channelrhodopsin-2 can be independent of intracortical glutamatergic synaptic transmission. PLoS One 7:1–10. doi:10.1371/journal.pone.0029859

Sharif NA, Klimko PG. 2019. Prostaglandin FP receptor antagonists: discovery, pharmacological characterization and therapeutic utility. Br J Pharmacol 176:1059–1078. doi:10.1111/bph.14335

Shmuel A, Yacoub E, Pfeuffer J, Moortele P Van De, Adriany G, Hu X, Ugurbil K, Van de Moortele PF, Adriany G, Hu X, Ugurbil K. 2002. Sustained negative BOLD, blood flow and oxygen consumption response and its coupling to the positive response in the human brain. Neuron 36:1195–1210. doi:10.1016/S0896-6273(02)01061-9

Smetters D, Majewska A, Yuste R. 1999. Detecting action potentials in neuronal populations with calcium imaging. Methods: A Companion to Methods in Enzymology 18:215–221. doi:10.1006/meth.1999.0774

Smith SJ, Smbül U, Graybuck LT, Collman F, Seshamani S, Gala R, Gliko O, Elabbady L, Miller JA, Bakken TE, Rossier J, Yao Z, Lein E, Zeng H, Tasic B, Hawrylycz M. 2019. Single-cell transcriptomic evidence for dense intracortical neuropeptide networks. Elife 8:1–35. doi:10.7554/eLife.47889

Sun W, McConnell E, Pare J-F, Xu Q, Chen M, Peng W, Lovatt D, Han X, Smith Y, Nedergaard M. 2013. Glutamate-Dependent Neuroglial Calcium Signaling Differs Between Young and Adult Brain. Science (1979) 339:197–200. doi:10.1126/science.1226740

Takei S, Hasegawa-Ishii S, Uekawa A, Chiba Y, Umegaki H, Hosokawa M, Woodward DF, Watanabe K, Shimada A. 2012. Immunohistochemical demonstration of increased prostaglandin F2α levels in the rat hippocampus following kainic acid-induced seizures. Neuroscience 218:295–304. doi:10.1016/j.neuroscience.2012.05.013

Takemiya T, Matsumura K, Yamagata K. 2007. Roles of prostaglandin synthesis in excitotoxic brain diseases. Neurochem Int 51:112–120. doi:10.1016/j.neuint.2007.05.009

Tasic B, Menon V, Nguyen TNTN, Kim TKTTKTK, Jarsky T, Yao Z, Levi BBP, Gray LT, Sorensen SA, Dolbeare T, Bertagnolli D, Goldy J, Shapovalova N, Parry S, Lee CCK, Smith K, Bernard A, Madisen L, Sunkin SM, Hawrylycz M, Koch C, Zeng H, Yao Z, Lee CCK, Shapovalova N, Parry S, Madisen L, Sunkin SM, Hawrylycz M, Koch C, Zeng H. 2016. Adult mouse cortical cell taxonomy revealed by single cell transcriptomics. Nat Neurosci advance on:1–37. doi:10.1038/nn.4216

Team RC, R Core Team, Team RC. 2023. R: A Language and Environment for Statistical Computing.

Thévenaz P, Ruttimann UE, Unser M. 1998. A pyramid approach to subpixel registration based on intensity. IEEE Transactions on Image Processing 7:27–41. doi:10.1109/83.650848

Tournissac M, Boido D, Omnès M, Goulam-Houssen Y, Ciobanu L, Charpak S. 2022. Cranial window for longitudinal and multimodal imaging of the whole mouse cortex. Neurophotonics 9. doi:10.1117/1.nph.9.3.031921

Tran CHT, George AG, Teskey GC, Gordon GR. 2020. Seizures elevate gliovascular unit Ca2+ and cause sustained vasoconstriction. JCI Insight 5. doi:10.1172/jci.insight.136469

Tsai CJ, Nagata T, Liu CY, Suganuma T, Kanda T, Miyazaki T, Liu K, Saitoh T, Nagase H, Lazarus M, Vogt KE, Yanagisawa M, Hayashi Y. 2021. Cerebral capillary blood flow upsurge during REM sleep is mediated by A2a receptors. Cell Rep 36. doi:10.1016/j.celrep.2021.109558

Tsurumaki T, Yamaguchi T, Higuchi H. 2002. Marked neuropeptide Y-induced contractions via NPY-Y1 receptor and its desensitization in rat veins. Vascul Pharmacol 39:325–333. doi:10.1016/S1537-1891(03)00044-2

Uhlirova H, Kılıç K, Tian P, Thunemann M, Desjardins M, Saisan PA, Sakadžić S, Ness T V., Mateo C, Cheng Q, Weldy KL, Razoux F, Vandenberghe M, Cremonesi JA, Ferri CGL, Nizar K, Sridhar VB, Steed TC, Abashin M, Fainman Y, Masliah E, Djurovic S, Andreassen OA, Silva GA, Boas DA, Kleinfeld D, Buxton RB, Einevol GT, Dale AM, Devor A, Kılıc K, Tian P, Thunemann M, Ness T V., Saisan PA, Sakadz S, Mateo C, Cheng Q, Weldy KL, Razoux F, Vandenberghe M, Cremonesi JA, Ferri CGL, Nizar K, Sridhar VB, Steed TC, Abashin M, Silva GA, Boas DA, Kleinfeld D, Buxton RB. 2016. Cell type specificity of neurovascular coupling in cerebral cortex. Elife 5:1–23. doi:10.7554/eLife.14315

Voutsinos-Porche B, Bonvento G, Tanaka K, Steiner P, Welker E, Chatton JY, Magistretti PJ, Pellerin L. 2003. Glial glutamate transporters mediate a functional metabolic crosstalk between neurons and astrocytes in the mouse developing cortex. Neuron 37:275–286. doi:10.1016/S0896-6273(02)01170-4

Wagner L, Wolf R, Zeitschel U, Rossner S, Petersén Å, Leavitt BR, Kästner F, Rothermundt M, Gärtner UT, Gündel D, Schlenzig D, Frerker N, Schade J, Manhart S, Rahfeld JU, Demuth HU, Von Hörsten S. 2015. Proteolytic degradation of neuropeptide y (NPY) from head to toe: Identification of novel NPY-cleaving peptidases and potential drug interactions in CNS and Periphery. J Neurochem 135:1019–1037. doi:10.1111/jnc.13378

Yamagata K, Andreasson KI, Kaufmann WE, Barnes CA, Worley PF. 1993. Expression of a Mitogen-Inducible Cyclooxygenase in Brain Neurons : Regulation by Synaptic Activity and Glucocorticoids. Neuron 11:371–386. doi:10.1016/0896-6273(93)90192-t

Yamagata K, Matsumura K, Inoue W, Shiraki T, Suzuki K, Yasuda S, Sugiura H, Cao C, Watanabe Y, Kobayashi S. 2001. Coexpression of microsomal-type prostaglandin E synthase with cyclooxygenase-2 in brain endothelial cells of rats during endotoxin-induced fever. Journal of Neuroscience 21:2669–2677. doi:10.1523/jneurosci.21-08-02669.2001

Zeisel A, Muñoz-Manchado AB, Codeluppi S, Lönnerberg P, La Manno G, Juréus A, Marques S, Munguba H, He L, Betsholtz C, Rolny C, Castelo-Branco G, Hjerling-Leffler J, Linnarsson S. 2015. Cell types in the mouse cortex and hippocampus revealed by single-cell RNA-seq. Science (1979) 347:1138–1142.

Zhang D, Ruan J, Peng S, Li J, Hu X, Zhang Y, Zhang T, Ge Y, Zhu Z, Xiao X, Zhu YY, Li XX, Li T, Zhou L, Gao Q, Zheng G, Zhao B, Li XX, Zhu YY, Wu J, Li W, Zhao J, Ge W, Xu T, Jia J-M. 2024. Synaptic-like transmission between neural axons and arteriolar smooth muscle cells drives cerebral neurovascular coupling. Nat Neurosci 1–17. doi:10.1038/s41593-023-01515-0

Zhang DY, Raichle ME. 2010. Disease and the brain’s dark energy. Nat Rev Neurol 6:15–28. doi:10.1038/nrneurol.2009.198

Zhou N, Gordon GRJ, Feighan D, MacVicar BA. 2010. Transient swelling, acidification, and mitochondrial depolarization occurs in neurons but not astrocytes during spreading depression. Cerebral Cortex 20:2614–2624. doi:10.1093/cercor/bhq018

Zonta M, Angulo MC, Gobbo S, Rosengarten B, Hossmann K-A, Pozzan T, Carmignoto G. 2003. Neuron-to-astrocyte signaling is central to the dynamic control of brain microcirculation. Nat Neurosci 6:43–50. doi:10.1038/nn980

