## Supplementary material for "Elevated pyramidal cell firing orchestrates arteriolar vasoconstriction through COX-2-derived prostaglandin E2 signaling": Supplmentary file

**Supplementary Informations**

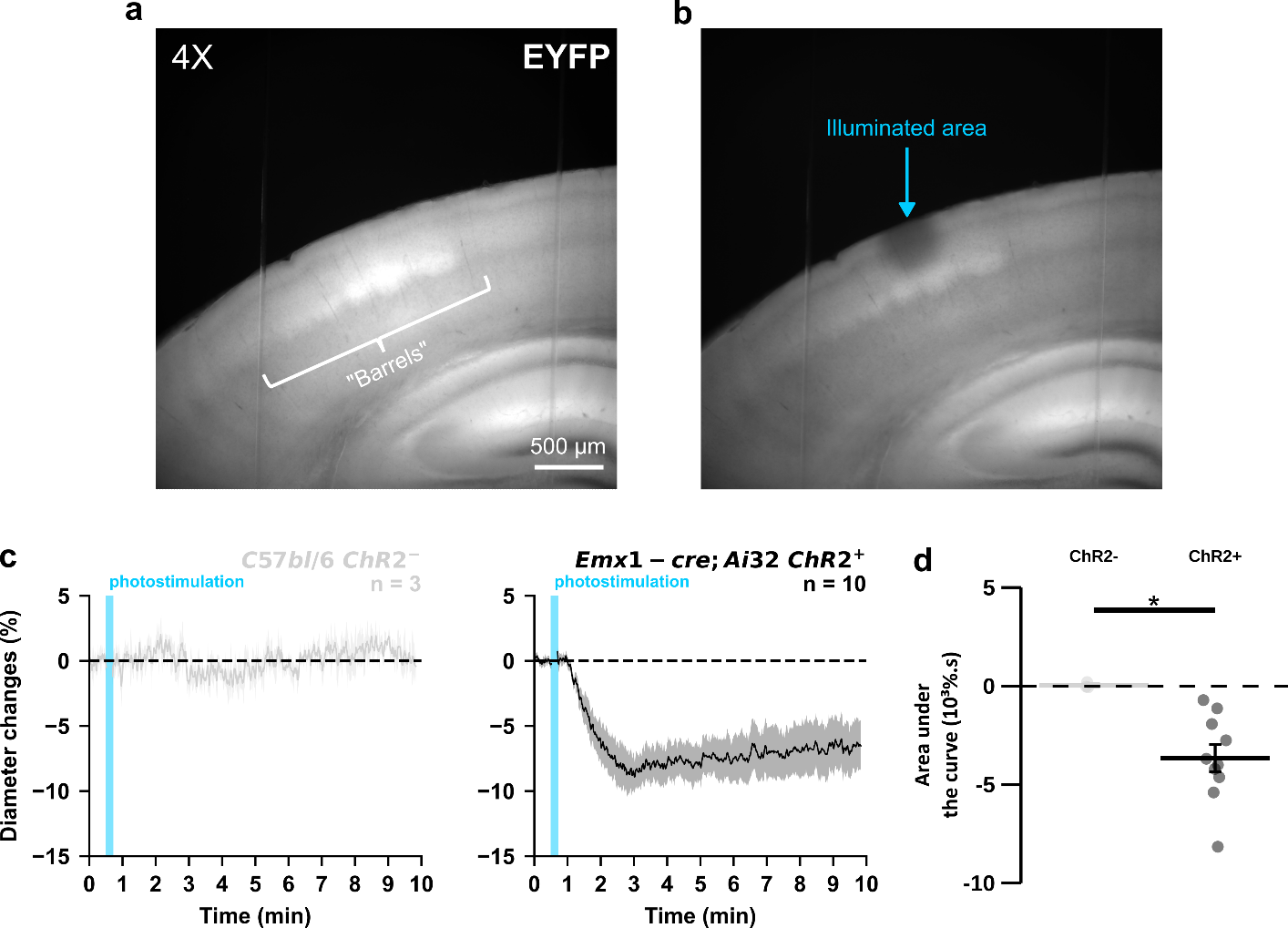

**Supplementary Figure 1: Vasoconstriction induced by widefield photostimulation is specific of ChR2 expression in pyramidal cells. (a)** Visualization of EYFP-ChR2 fusion transgene fluorescence in a cortical slice of an Emx1-cre;Ai32 mouse at 4X objective. Note the presence of barrels in layer IV. **(b)** Photobleaching in superficial cortical layers was achieved by widefield illumination at maximum LED power for one minute with a 40X objective. The round photobleached area was approximately 0.15 mm². **(c)** Kinetics of vascular responses induced by photostimulation at 20 Hz in cortical brain slices from naive C57bl/6J (light gray, n= 3 arterioles) or ChR2-expressing Emx1-cre;Ai32 mice (black, n= 10 arterioles). Dashed line represents the baseline. The SEMs envelope the mean traces. **(d)** Effect of pyramidal cell ChR2 expression on AUC of vascular responses evoked by photostimulation at 20 Hz. The data are shown as the individual values and mean ± SEM. * statistically different with p< 0.05.

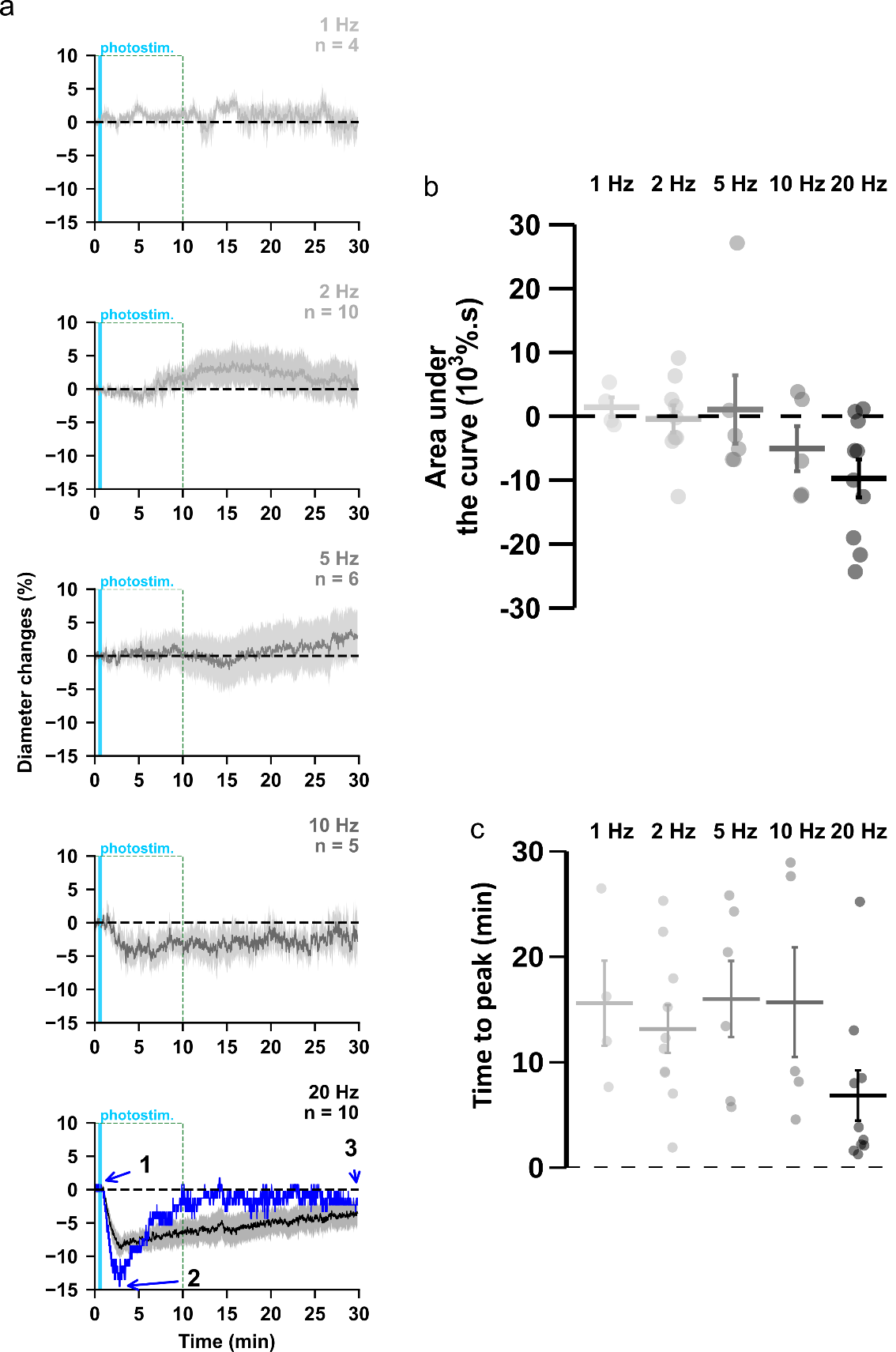

**Supplementary Figure 2: Vasoconstrictions occurred during the 10 first minutes after pyramidal cells photoactivation.** **(a)** Kinetics of arteriolar diameter changes induced by photostimulation (vertical cyan bars) at 1 Hz (n= 4 arterioles from 3 mice), 2 Hz (n= 10 arterioles from 8 mice), 5 Hz (n= 6 arterioles from 6 mice), 10 Hz (n= 5 arterioles from 5 mice) and 20 Hz (n= 10 arterioles from 9 mice) during 30 min recording. The SEMs envelope the mean traces. The blue trace represents the kinetics of the diameter changes of the arteriole shown in Fig. 1b. **(b-c)** Effects of the different photostimulation frequencies on (b) AUC and (c) of vascular responses during the 30 min of recording. Data are presented as the individual values and mean ± SEM.

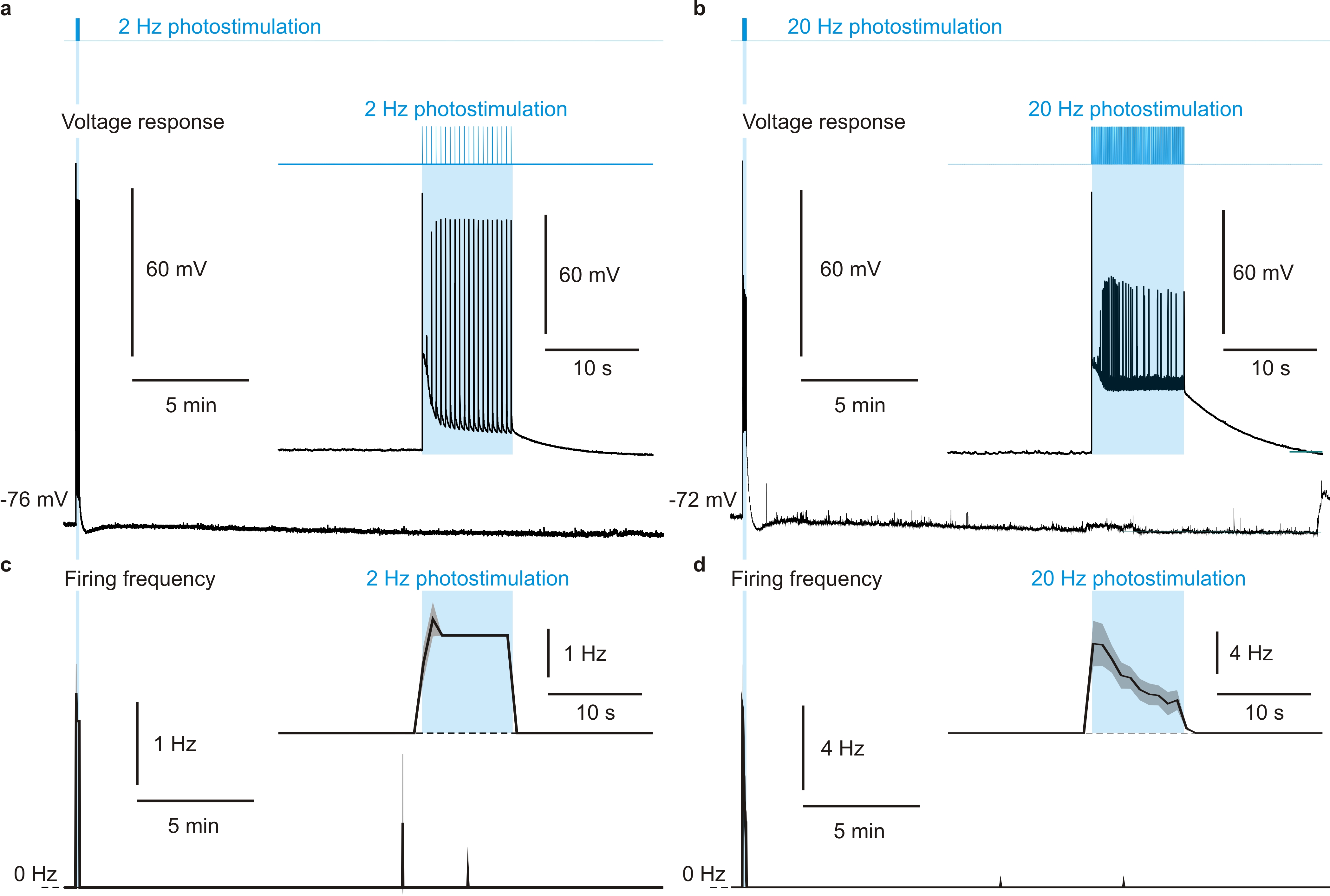

**Supplementary Figure 3: Photostimulation of pyramidal cells does not evoke recurrent spiking network activity. (a,b)** Representative voltage responses evoked before, during and after photostimulation (vertical cyan bars) delivered at 2 (a) and 20 Hz (b). The insets show the enlarged voltage responses during photostimulation (cyan zones). **(c,d)** Mean firing frequency of pyramidal cells evoked at 2 Hz (**c**, n= 9 cells) and 20 Hz (**d**, n= 9 cells). Note sporadic action potentials after photostimulation. The insets show the zoomed-in mean firing frequency during the 40-s period around photostimulation. The dashed line represents the 0 Hz baseline. The SEMs envelope the mean traces.

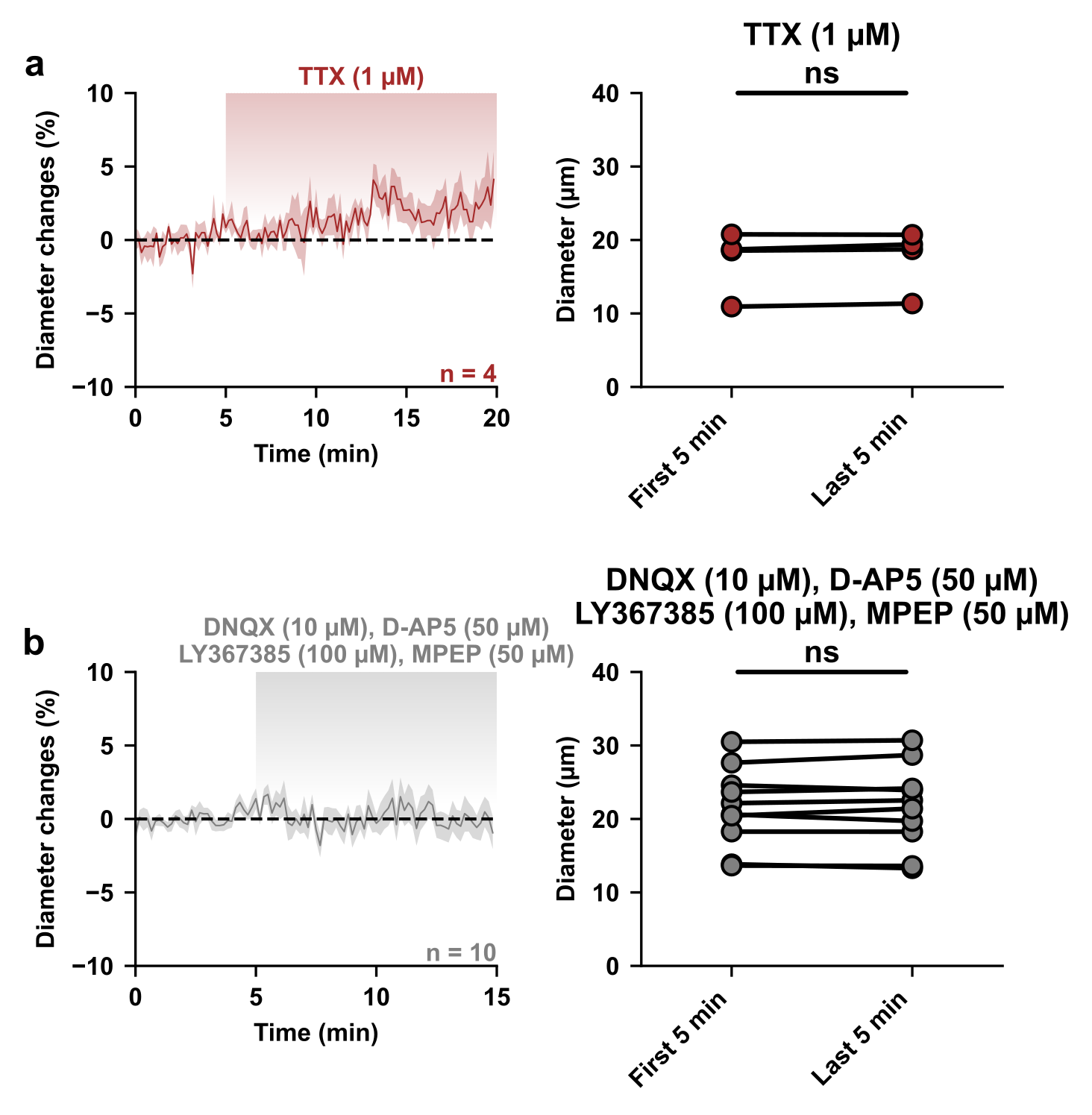

**Supplementary Figure 4: Basal network activity and tonic glutamate do not influence resting vascular tone.** **(a,b)** Kinetics of diameter changes (left panels) and comparison of the mean luminal diameter between the five-minute control period and after 15 or 20 minutes of treatment (right panels) with (**a**) TTX (1 µM, brown, n= 4 arterioles) or (**b**) a cocktail of AMPA/kainate (DNQX, 10 µM), NMDA (D-AP5, 50 µM), mGluR1 (LY367385, 100 µM) and mGluR5 (MPEP, 50 µM) glutamate receptor antagonists (gray, n= 10 arterioles). n.s. not statistically significant.

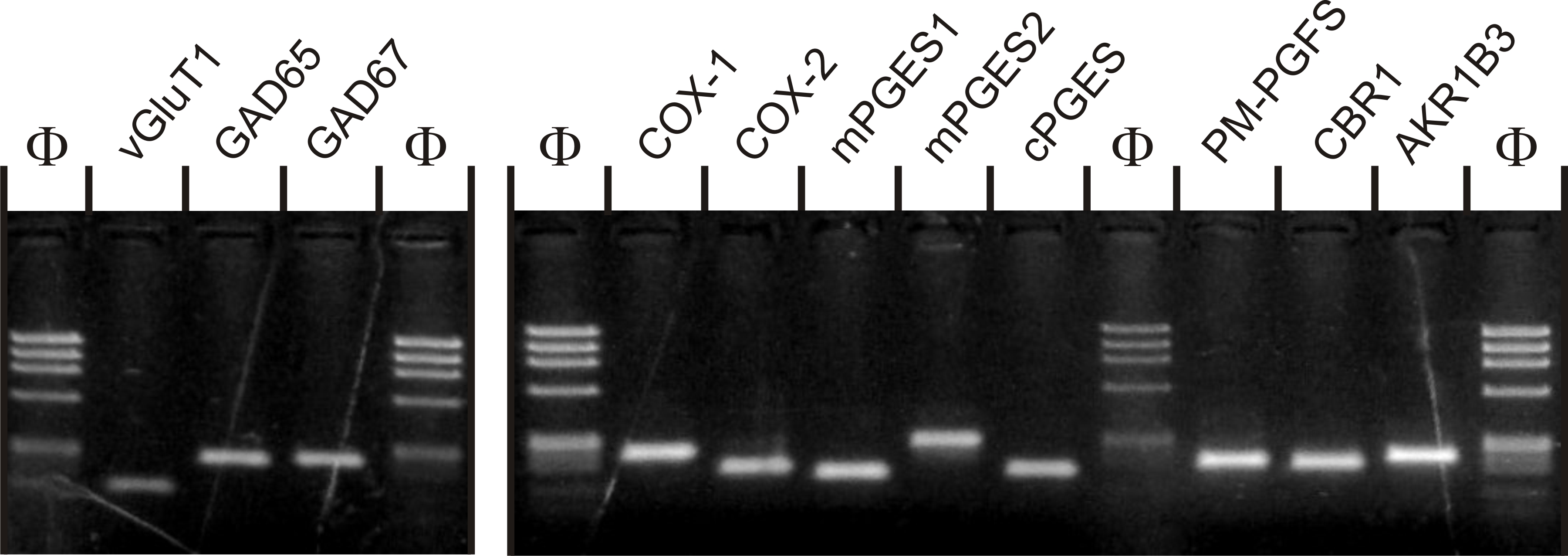

**Supplementary Figure 5: Sensitivity of the RT-mPCR protocol**. Agarose gel analysis of a RT-PCR performed from 500 pg of forebrain total RNAs Φx174 digested by *HaeIII* (Φ) was used as molecular weight marker. All the amplicons were detected with the expected sized from the gene sequences (supplementary Table 2).

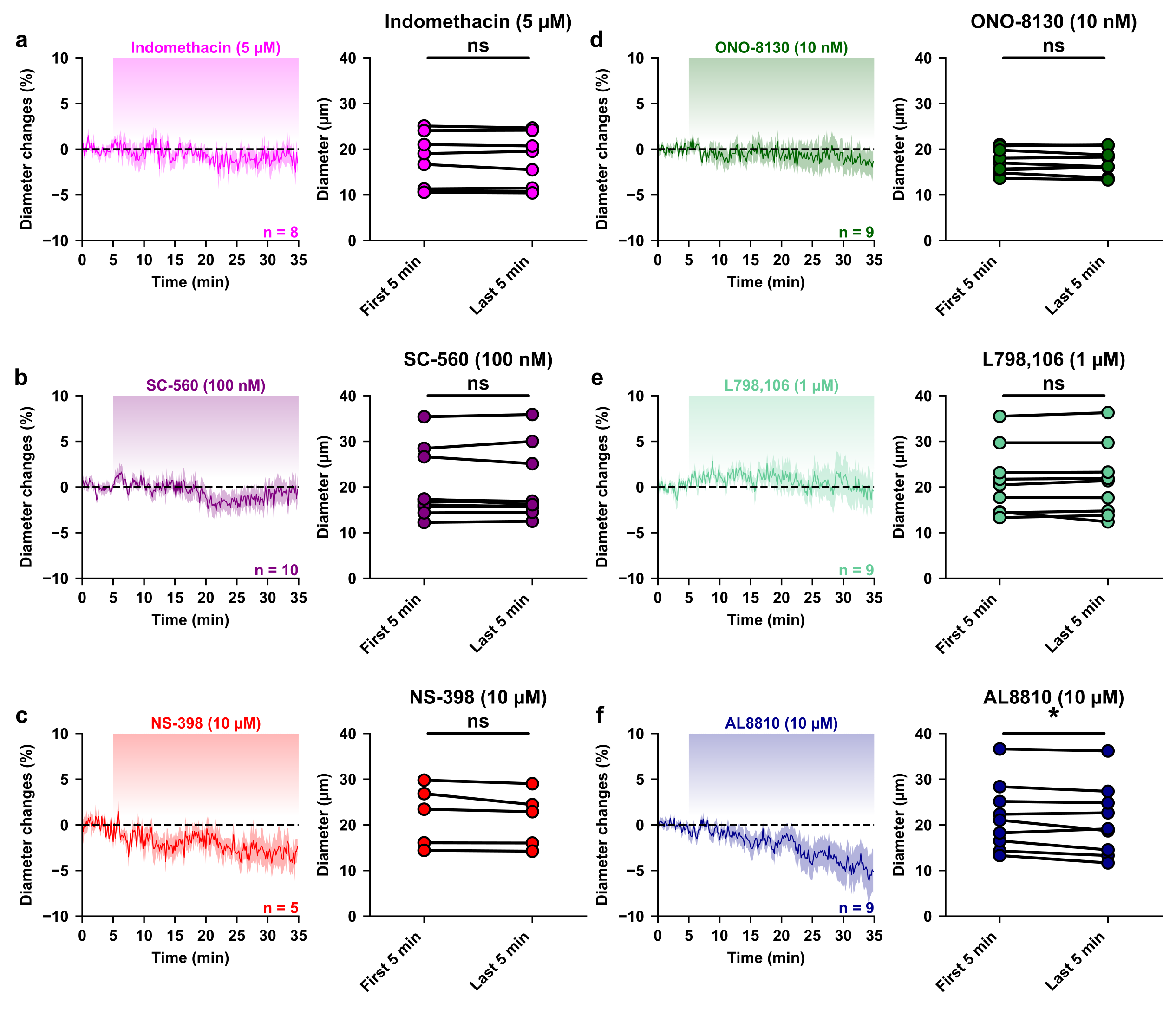

**Supplementary Figure 6: Tonic PGE2 does not affect basal vascular tone.** Kinetics of diameter changes (left panels) and comparison of the mean luminal diameter between the five minutes control period and the five minutes following 30 minutes treatment by the COX inhibitors **(a)** indomethacin (5 µM, magenta, n= 8 arterioles), **(b)** SC-560 (100 nM, purple, n= 10 arterioles), **(c)** NS-398 (10 µM, red, n= 5 arterioles), and the EP1, EP3 and FP antagonists **(d)** ONO-8130 (10 nM, dark green, n= 9 arterioles), **(e)** L-798,106 (1 µM, light green, n= 9 arterioles) and **(f)** AL8810 (10 µM, dark blue, n= 9 arterioles), respectively. n.s. not statistically significant and * statistically different with p< 0.05.

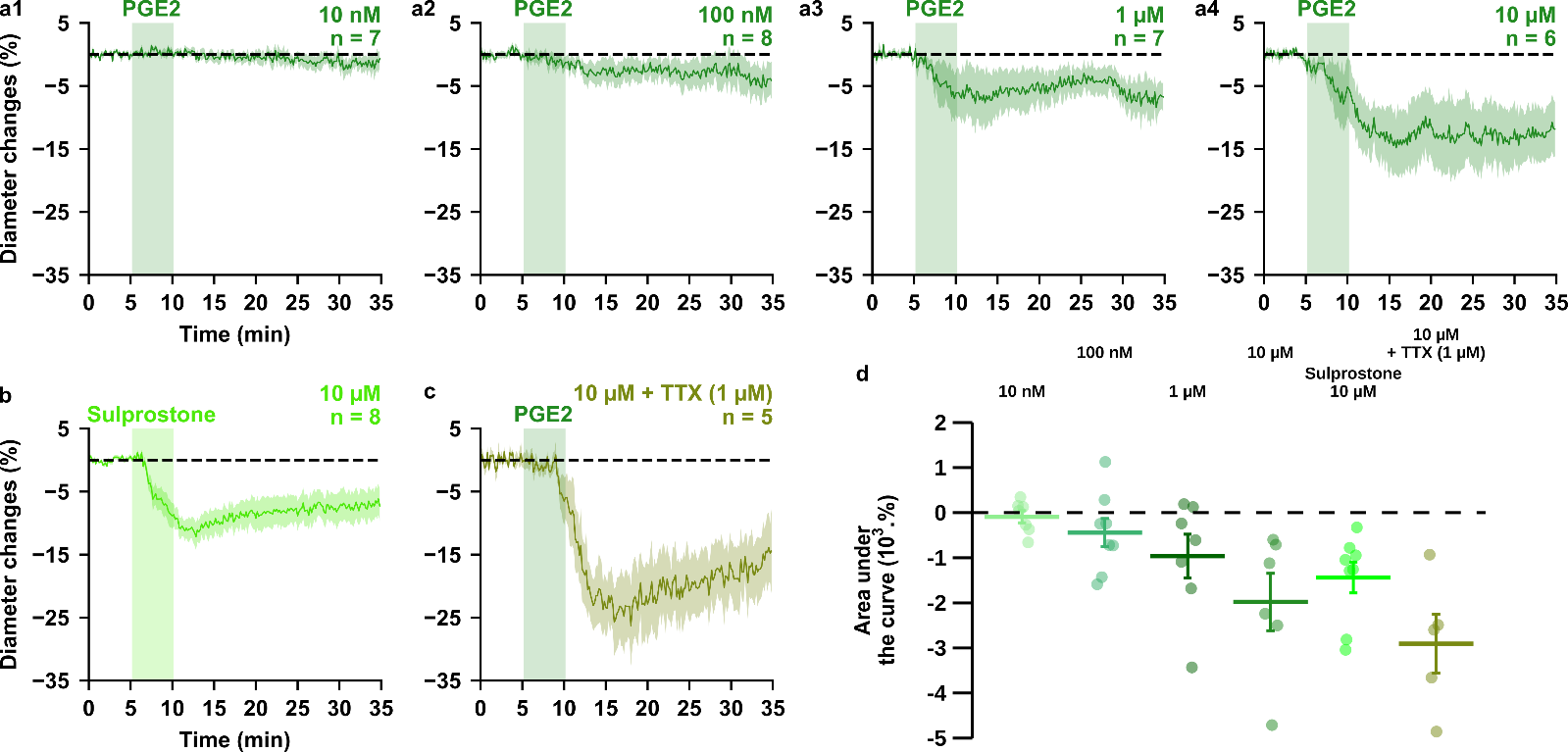

**Supplementary Figure 7: PGE2 dose-dependently induces vasoconstriction. (a, b)** Kinetics of arteriolar diameter changes induced by exogenous application of PGE2 (vertical green zones) at **(a1)** 10 nM (n= 7 arterioles from 4 mice), **(a2)** 100 nM (n=8 arterioles from 6 mice), **(a3)** 1 µM (n= 7 arterioles from 4 mice) and **(a4)** 10 µM (n= 6 arterioles from 4 mice), by **(b)** the EP1/EP3 agonist sulprostone (fluorescent green, 10 µM, n= 8 arterioles from 5 mice) or **(c)** by 10 µM PGE2 under a TTX application (1 µM). Dashed line represents the baseline. The SEMs envelope the mean traces. **(d)** Dose-dependent effect of PGE2 or sulprostone effect on AUC of vascular responses. The data are shown as the individual values and mean ± SEM.

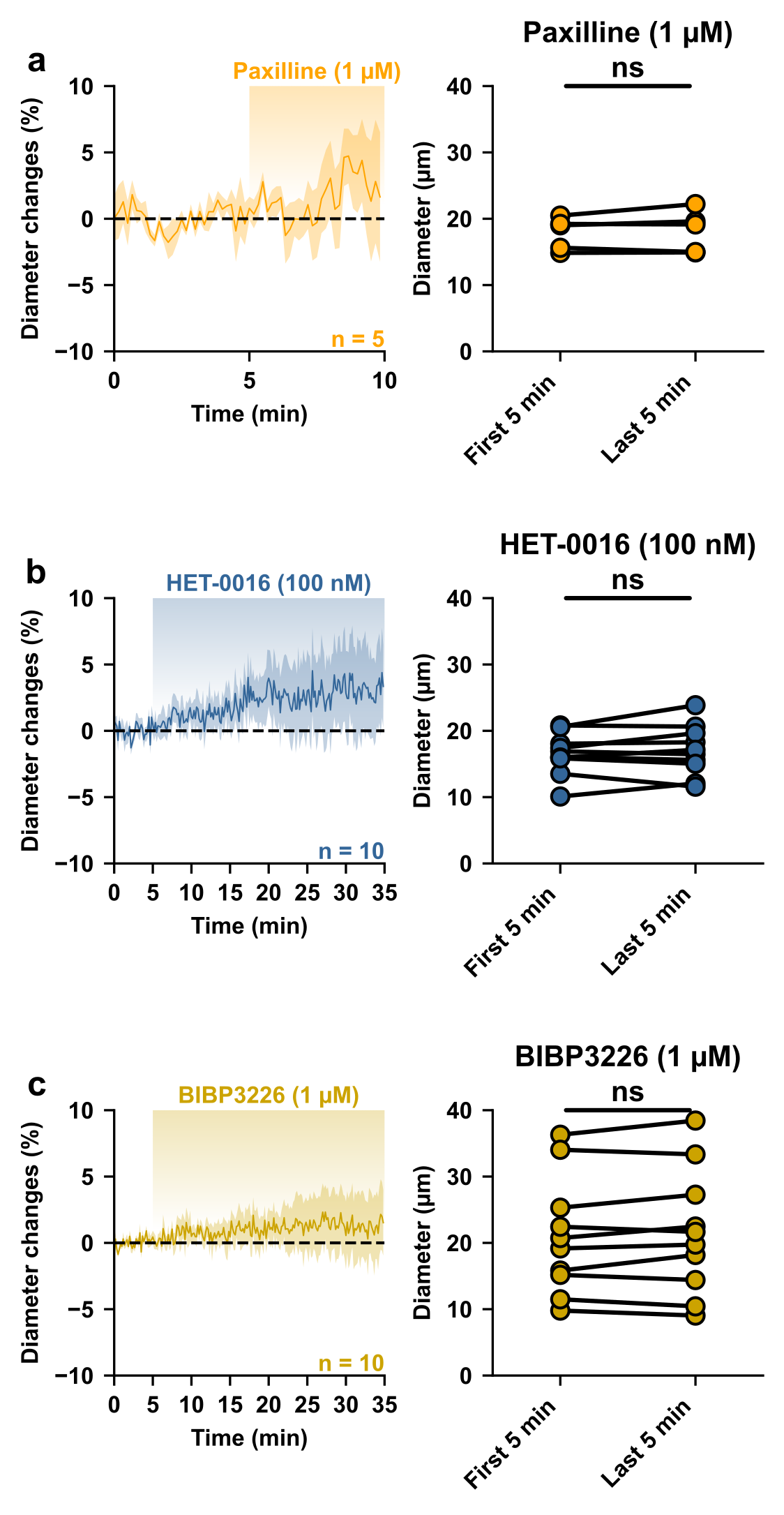

**Supplementary Figure 8: Vasoconstrictive pathways do not influence resting vascular tone.** Kinetics of diameter changes (left panels) and comparison of the mean luminal diameter between the five-minute control period and the last five minutes (right panels) of treatment with paxilline for five minutes (**a,** 1 µM, orange, n= 5 arterioles) or with HET-0016 (**b,** 100 nM, blue-grey, n= 10 arterioles) or BIBP3226 (**c**, 1 µM, yellow, n=10 arterioles) for 30 minutes. n.s. not statistically significant.

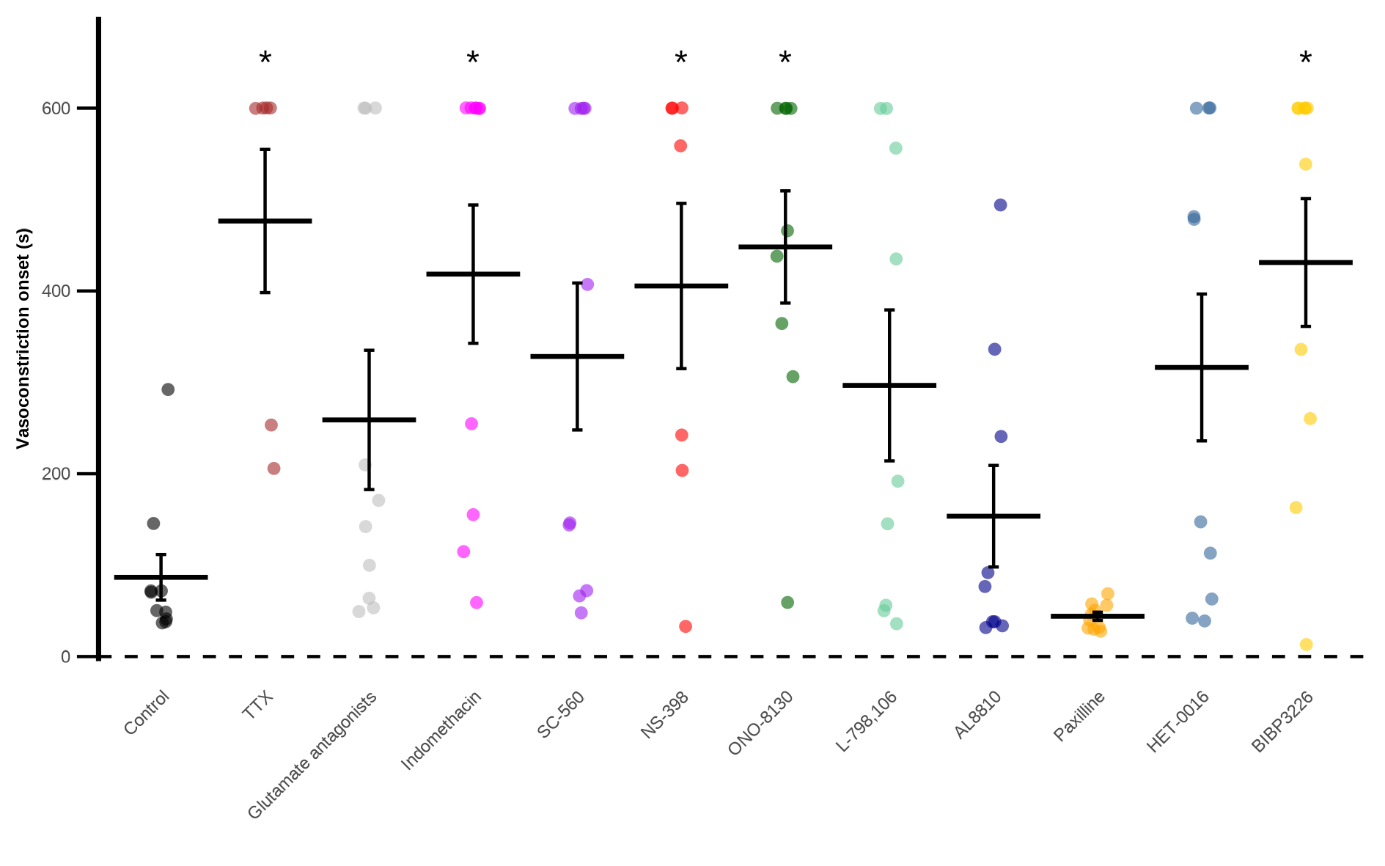

**Supplementary Figure 9: Vasoconstriction onset times across the different conditions.** Onset times of vasoconstriction after a 20 Hz photostimulation (measured in seconds after the beginning of the photostimulation) in control, TTX (1 µM, brown, n= 4 arterioles), a cocktail of glutamate antagonists (AMPA/kainate (DNQX, 10 µM), NMDA (D-AP5, 50 µM), mGluR1 (LY367385, 100 µM) and mGluR5 (MPEP, 50 µM) antagonists; gray, n= 10 arterioles), indomethacin (5 µM, magenta, n= 8 arterioles), SC-560 (100 nM, purple, n= 10 arterioles), NS-398 (10 µM, red, n= 5 arterioles), ONO-8130 (10 nM, dark green, n= 9 arterioles), L-798,106 (1 µM, light green, n= 9 arterioles), AL8810 (10 µM, dark blue, n= 9 arterioles), Paxilline (1 µM, orange, n= 10 arterioles), HET-0016(100 nM, blue-grey, n= 10 arterioles), and BIBP3226 (1 µM, yellow, n=10 arterioles) condition. The data are shown as the individual values and mean ± SEM. * statistically different from control with p< 0.05.

**Supplementary table 1:** **Morphological and physiological properties, and neurovascular responses of diving arterioles used in the analysis of the frequency-dependence of the polarity of neurovascular response evoked by pyramidal cells.**

| **Frequency** | **1 Hz** | **2 Hz** | **5 Hz** | **10 Hz** | **20 Hz** |
| --- | --- | --- | --- | --- | --- |
| Number of arterioles | n = 4 | n = 10 | n = 6 | n = 5 | n = 10 |
| **Resting stability (%)** | 1.2 ± 0.2 | 1.6 ± 0.2 | 1.6 ± 0.2 | 1.3 ± 0.2 | 1.0 ± 0.1 |
|  | F (4, 30) = 2.161  p = 0.098 | | | | |
|  | n.s. | | | | |
| **Wall thickness (µm)** | 3.6 ± 0.8 | 3.8 ± 0.3 | 3.2 ± 0.5 | 4.1 ± 0.8 | 4.0 ± 0.2 |
|  | F (4, 30) = 0.656  p = 0.627 | | | | |
|  | n.s. | | | | |
| **Area under the curve after photostimulation (AUC; x10^3^ %.s)** | 0.5 ± 0.2 | 0.0 ± 0.5 | 0.2 ± 1 | -1.7 ± 1.1 | -3.7 ± 0.7 |
|  | F (4, 30) = 6.135  p = 0.00099 | | | | |
|  | *** | | | | |
| **Maximal dΔT/dt (%.s^-1^)** | 0.19 ± 0.09 | 0.64 ± 0.14 | 0.22 ± 0.04 | 0.43 ± 0.19 | 0.8 ± 0.11 |
|  | All < 2 %.s^-1^ | | | | |

Data are mean ± SEM, one-way ANOVA F test and corresponding exact p-value. n.s., not statistically different and ***: p<0.001.

**Supplementary table 2:** PCR primers

| **Gene accession #** | **First PCR primers** | **Size (bp)** | **Second PCR nested primers** | **Size (bp)** |
| --- | --- | --- | --- | --- |
| vGluT1 | Sense, -113: GGCTCCTTTTTCTGGGGCTAC | 259 | Sense, -54: ATTCGCAGCCAACAGGGTCT | 153 |
| NM_182993 | (Cabezas et al., 2013) |  | (Cabezas et al., 2013) |  |
|  | Antisense, 126: CCAGCCGACTCCGTTCTAAG |  | Antisense, 79: TGGCAAGCAGGGTATGTGAC |  |
|  | (Gallopin et al., 2006) |  | (Cabezas et al., 2013) |  |
| GAD65 | Sense, 99: CCAAAAGTTCACGGGCGG | 375 | Sense, 219: CACCTGCGACCAAAAACCCT | 248 |
| NM_008078 | (Karagiannis et al., 2009) |  | (Perrenoud et al., 2012) |  |
|  | Antisense, 454: TCCTCCAGATTTTGCGGTTG |  | Antisense, 447: GATTTTGCGGTTGGTCTGCC |  |
|  | (Karagiannis et al., 2009) |  | (Perrenoud et al., 2012) |  |
| GAD67 | Sense, 529: TACGGGGTTCGCACAGGTC | 598 | Sense, 801: CCCAGAAGTGAAGACAAAAGGC | 255 |
| NM_008077 | (Férézou et al., 2002) |  | (Cabezas et al., 2013) |  |
|  | Antisense, 1,109: CCCAGGCAGCATCCACAT |  | Antisense, 1,034: AATGCTCCGTAAACAGTCGTGC |  |
|  | (Cabezas et al., 2013) |  | (Cabezas et al., 2013) |  |
| COX1 | Sense, 107: ATCCCTGTTGTTACTATCCGTGC | 383 | Sense, 137: AGGGTGTCTGTGTCCGCTTT | 249 |
| NM_008969 | Antisense, 470: TGTGGGGCAGTCTTTGGGTA |  | Antisense, 366: GGCTGGGGATAAGGTTGGAC |  |
| COX2 | Sense, 199: CTGAAGCCCACCCCAAACAC | 268 | Sense, 265: AACAACATCCCCTTCCTGCG | 181 |
| NM_011198 | (Lecrux et al., 2011) |  | (Devienne et al., 2018) |  |
|  | Antisense, 445: CCTTATTTCCCTTCACACCCAT |  | Antisense, 426: TGGGAGTTGGGCAGTCATCT |  |
|  | (Devienne et al., 2018) |  | (Devienne et al., 2018) |  |
| mPGES1 | Sense, 14: GCCTGGTGATGGAGAGCG | 371 | Sense, 110: AGATGAGGCTGCGGAAGAAG | 158 |
| NM_022415 | Antisense, 367: GGAGCGAAGGCGTGGGTT |  | Antisense, 248: CACGAAGCCGAGGAAGAGGA |  |
| mPGES2 | Sense, 357: CGACTTCCACTCCCTGCC | 337 | Sense, 388: GAGGTGAATCCCGTGAGAAGG | 288 |
| NM_133783 | Antisense, 374: CATCTCCTCCGTCCTGGCTT |  | Antisense, 656: TTCCTTCCCGCCATACATCT |  |
| cPGES | Sense, 190: TCCAAGCATAAAAGAACAGACAGA | 282 | Sense, 266: TAACAAAGGAAAGGGCAAAGC | 174 |
| NM_019766 | Antisense, 448: TGGCATCTTTTCATCATCACTGTC |  | Antisense, 416: CATCATCTGCTCCATCTACTTCTG |  |
| PM-PGFS | Sense, 236: AGGAGTTTCTGGATGGTGGTTAC | 370 | Sense, 360: ACCTGTTCGTGATGTAGCCTCC | 220 |
| NM_025582 | Antisense, 584: CACCTCCCACACACCTCTTCAT |  | Antisense, 562: CTGGGGTGGCTTGCTGGA |  |
| Akr1b3 | Sense, 148: CAGAATGAGAAGGAGGTGGGA | 341 | Sense, 189: CAAGGAGCAGGTGGTGAAGC | 247 |
| NM_009658 | Antisense, 467: TTGAAGTTGGAGACACCGATTG |  | Antisense, 416: CATAGCCGTCCAAGTGTCCA |  |
| CBR1 | Sense, 196: AACCCGCAGAGCATTCGC | 373 | Sense, 291: CAATGACGACACCCCCTTCC | 208 |
| NM_007620 | Antisense, 551: GCCAACCTTCTTCCGCAT |  | Antisense, 477: CTCCTCTGTGATGGTCTCGCTT |  |

**Supplementary table 3:** **Electrophysiological properties of pyramidal cells recorded during single cell RT-PCR experiments.**

|  | **COXs-negative**  **(n=7)** | **COX-1 positive**  **(n=4)** | **COX-2 positive**  **(n=5)** |
| --- | --- | --- | --- |
| **Passive properties** | | |  |
| **(1) Resting potential (mV)** | -82.0 ± 2.2 | -84.7 ± 4.2 | -82.0 ± 6.1 |
| **(2) Input resistance (MΩ)** | 329 ± 53.7 | 360.8 ± 63.7 | 314.2 ± 79.7 |
| **(3) Time constant (ms)** | 50.7 ± 6.4 | 47.3 ± 7.2 | 47.74 ± 10.1 |
| **(4) Membrane capacitance (pF)** | 161.4 ± 14.7 | 133.2 ± 7.8 | 159.4 ± 23.0 |
| **(5) Sag index (%)** | 11.3 ± 3.8 | 6.7 ± 1.1 | 6.9 ± 1.8 |
| **Just above threshold properties** | | |  |
| **(6) Rheobase (pA)** | 52.7 ± 8.7 | 51.7 ± 15.5 | 62.3 ± 18.2 |
| **(7) First spike latency (ms)** | 295.2 ± 50.8 | 271.6 ± 62.9 | 178.7 ± 55.6 |
| **(8) Adaptation (Hz/s)** | -3.1 ± 0.9 | -2.6 ± 0.3 | -3.3 ± 1.3 |
| **(9) Minimal frequency (Hz)** | 5.5 ± 0.7 | 4.6 ± 0.4 | 6.3 ± 1.2 |
| **Firing properties** | | |  |
| **(10) Accommodation (mV)** | 16.4 ± 4.6 | 24.8 ± 4.9 | 11.9 ± 3.7 |
| **(11) Amplitude of early adaptation (Hz)** | 62.1 ± 13.3 | 89.8 ± 8.5 | 62.3 ± 16.4 |
| **(12) Time constant of early adaptation (ms)** | 29.7 ± 4.4 | 28.3 ± 2.2 | 43.1 ± 18.7 |
| **(13) Late adaptation (Hz/s)** | -11.6 ± 2.3 | -9.6 ± 3.9 | -10.7 ± 1.7 |
| **(14) Maximal frequency (Hz)** | 23.4 ± 1.7 | 27.6 ± 3.3 | 22.0 ± 2.9 |
| **Action potentials properties** | | |  |
| **(15) 1^st^ spike amplitude (mV)** | 94.8 ± 1.7 | 93.0 ± 5.0 | 88.6 ± 1.9 |
| **(16) 1^st^ spike duration (ms)** | 1.8 ± 0.1 | 1.9 ± 0.1 | 1.8 ± 0.1 |
| **(17) 2^nd^ spike amplitude (mV)** | 91.6 ± 1.9 | 91.9 ± 4.5 | 85.5 ± 2.5 |
| **(18) 2^nd^ spike duration (ms)** | 1.9 ± 0.1 | 1.9 ± 0.1 | 1.9 ± 0.1 |
| **(19) Amplitude Reduction (%)** | 3.4 ± 0.4 | 1.1 ± 0.7 | 3.6 ± 1.4 |
| **(20) Duration Increase (%)** | 5.3 ± 0.9 | 3.8 ± 1.5 | 7.5 ± 1.6 |
| **AHP and ADP properties** | | |  |
| **(21) 1^st^ spike fast AHP (mV)** | -8.7± 0.8 | -8.7 ± 0.6 | -7.9 ± 1.1 |
| **(22) 1^st^ spike ADP (mV)** | 0.2 ± 0.1 | 0.2 ± 0.2 | 0 ± 0 |
| **(23) 1^st^ spike medium AHP (mV)** | -13.8 ± 1.4 | -15.5 ± 0.6 | -13.7 ± 0.8 |
| **(24) 1^st^ spike fast AHP latency (ms)** | 8.2 ± 1.0 | 7.7 ± 1.4 | 9.8 ± 1.5 |
| **(25) 1^st^ spike ADP latency (ms)** | 4.6 ± 2.2 | 2.0 ± 2.0 | 0 ± 0 |
| **(26) 1^st^ spike, medium AHP latency (ms)** | 49.2 ± 3.3 | 48.2 ± 5.4 | 54.8 ± 6.1 |
| **(27) 2^nd^ spike fast AHP (mV)** | -9.8 ± 1.1 | -8.5 ± 0.5 | -9.6 ± 1.0 |
| **(28) 2^nd^ spike ADP (mV)** | 0 ± 0 | 0.1 ± 0.1 | 0 ± 0 |
| **(29) 2^nd^ spike medium AHP (mV)** | -16.1 ± 1.2 | -17.1 ± 0.6 | -15.8 ± 0.4 |
| **(30) 2^nd^ spike, fast AHP latency (ms)** | 8.6 ± 0.9 | 7.1 ± 0.7 | 11.7 ± 1.3 |
|  | F (2.13) = 4.063  p = 0.0426 | | |
|  | * | | |
|  | No significant difference in multiple comparisons. | | |
| **(31) 2^nd^ spike ADP latency (ms)** | 1.2 ± 1.2 | 1.7 ± 1.7 | 0 ± 0 |
| **(32) 2^nd^ spike, medium AHP latency (ms)** | 57.1 ± 5.8 | 51.9 ± 4.6 | 58.3 ± 6.7 |

Data are presented as mean ± SEM. Statistical analyses were performed using a one-way ANOVA (F test) or a Kruskal-Wallis test, depending on the result of the Shapiro-Wilk normality test. If a significant result was found, the corresponding statistics are reported, and post-hoc multiple comparisons are performed.

**Supplementary table 4: IC_50_ and concentrations of inhibitors, blocker and antagonists used in tissue.**

|  |  |  | **Concentrations used for inhibition/antagonism** | |
| --- | --- | --- | --- | --- |
| **Inhibitor/antagonist** | **In vitro IC_50_** |  | **Preparation** | **Concentration** |
| Indomethacin | COX-1: 22 nM  (Lora et al., 1998) | COX-2: 87 nM | Mouse brain slices  (Lacroix et al., 2015) | 5 µM |
| SC560 | COX-1: 9 nM  (Smith et al., 1998) | COX-2: 6.3 µM | Mouse brain slices  (Lacroix et al., 2015) | 100 nM |
| NS-398 | COX-1: 50 µM  (Lora et al., 1998) | COX-2: 0.6 µM | Mouse brain slices  (Lacroix et al., 2015) | 10 µM |
| ONO-8130 | EP1 receptors: 9.3 nM  (Säfholm et al., 2013b) |  | isolated guinea pig trachea  (Säfholm et al., 2013a) | 10 nM |
| L798,106 | EP3 receptors: 0.3 nM (Ki)  (Juteau et al., 2001) |  | Isolated mouse mesenteric arteries  (Chia et al., 2011) | 1 µM |
| AL8810 | FP receptors: 426 nM (Ki)  (Griffin et al., 1999) |  | Isolated porcine retinal arterioles  (Hansen et al., 2015) | 10 µM |
| Paxilline | BK channels: 97 nM  (Tammaro et al., 2004) |  | Mouse brain slices  (Girouard et al., 2010) | 1 µM |
| HET-0016 | CYP4A isoforms: 35 nM  (Miyata et al., 2001) |  | Mouse brain slices  (Blanco et al., 2008) | 100 nM |
| BIBP3226 | Y1 receptors: 26 nM  (Rudolf et al., 1994) |  | Mouse brain slices  (Bacci et al., 2002; Sun et al., 2003) | 1 µM |

**Supplementary References**

Bacci A, Huguenard JR, Prince D a. 2002. Differential modulation of synaptic transmission by neuropeptide Y in rat neocortical neurons. *Proc Natl Acad Sci U S A* **99**:17125–30. doi:10.1073/pnas.012481899

Blanco M, Stern JE, Filosa JA. 2008. Tone-dependent vascular responses to astrocyte-derived signals. *American journal of physiology* 2855–2863. doi:10.1152/ajpheart.91451.2007.

Cabezas C, Irinopoulou T, Cauli B, Poncer JC. 2013. Molecular and functional characterization of GAD67-expressing, newborn granule cells in mouse dentate gyrus. *Front Neural Circuits* **7**:1–16. doi:10.3389/fncir.2013.00060

Chia E, Kagota S, Wijekoon EP, McGuire JJ. 2011. Protection of protease-activated receptor 2 mediated vasodilatation against angiotensin II-induced vascular dysfunction in mice. *BMC Pharmacol* **11**. doi:10.1186/1471-2210-11-10

Devienne G, Le Gac B, Piquet J, Cauli B. 2018. Single Cell Multiplex Reverse Transcription Polymerase Chain Reaction After Patch-Clamp. *Journal of Visualized Experiments* **136**:1–12. doi:10.3791/57627

Férézou I, Cauli B, Hill EL, Rossier J, Hamel E, Lambolez B. 2002. 5-HT3 receptors mediate serotonergic fast synaptic excitation of neocortical vasoactive intestinal peptide/cholecystokinin interneurons. *Journal of Neuroscience* **22**:7389–7397. doi:10.1523/jneurosci.22-17-07389.2002

Gallopin T, Geoffroy H, Rossier J, Lambolez B. 2006. Cortical sources of CRF, NKB, and CCK and their effects on pyramidal cells in the neocortex. *Cerebral Cortex* **16**:1440–1452. doi:10.1093/cercor/bhj081

Girouard H, Bonev AD, Hannah RM, Meredith A, Aldrich RW, Nelson MT. 2010. Astrocytic endfoot Ca2+ and BK channels determine both arteriolar dilation and constriction. *Proc Natl Acad Sci U S A* **107**:3811–3816. doi:10.1073/pnas.0914722107

Griffin BW, Klimko P, Crider JY, Sharif NA. 1999. AL-8810: A novel prostaglandin F(2??) analog with selective antagonist effects at the prostaglandin F(2??) (FP) receptor. *Journal of Pharmacology and Experimental Therapeutics* **290**:1278–1284.

Hansen PO, Kringelholt S, Simonsen U, Bek T. 2015. Hypoxia-induced relaxation of porcine retinal arterioles in vitro depends on inducible NO synthase and EP 4 receptor stimulation in the perivascular retina 457–463. doi:10.1111/aos.12669

Juteau H, Gareau Y, Labelle M, Lamontagne S, Tremblay N, Carrière MC, Sawyer N, Denis D, Metters KM. 2001. Structure-activity relationship on the human EP3 prostanoid receptor by use of solid-support chemistry. *Bioorg Med Chem Lett* **11**:747–749. doi:10.1016/S0960-894X(01)00056-7

Karagiannis A, Gallopin T, Dávid C, Battaglia D, Geoffroy H, Rossier J, Hillman EMC, Staiger JF, Cauli B. 2009. Classification of NPY-expressing neocortical interneurons. *J Neurosci* **29**:3642–59. doi:10.1523/JNEUROSCI.0058-09.2009

Lacroix A, Toussay X, Anenberg E, Lecrux C, Ferreirós N, Karagiannis A, Plaisier F, Chausson P, Jarlier F, Burgess SA, Hillman EMCCC, Tegeder I, Murphy TH, Hamel E, Cauli B, Ferreiro N, Burgess SA, Hillman EMCCC, Tegeder I, Murphy TH, Hamel E, Cauli B. 2015. COX-2-Derived Prostaglandin E2 Produced by Pyramidal Neurons Contributes to Neurovascular Coupling in the Rodent Cerebral Cortex. *J Neurosci* **35**:11791–11810. doi:10.1523/JNEUROSCI.0651-15.2015

Lecrux C, Toussay X, Kocharyan A, Fernandes P, Neupane S, Levesque M, Plaisier F, Shmuel A, Cauli B, Hamel E. 2011. Pyramidal Neurons Are “Neurogenic Hubs” in the Neurovascular Coupling Response to Whisker Stimulation. *Journal of Neuroscience* **31**:9836–9847. doi:10.1523/JNEUROSCI.4943-10.2011

Lora M, Denault JB, Leduc R, De Brum-Fernandes AJ. 1998. Systematic pharmacological approach to the characterization of NSAIDs. *Prostaglandins Leukot Essent Fatty Acids* **59**:55–62. doi:10.1016/S0952-3278(98)90052-7

Miyata N, Taniguchi K, Seki T, Ishimoto T, Sato-Watanabe M, Yasuda Y, Doi M, Kametani S, Tomishima Y, Ueki T, Sato M, Kameo K. 2001. HET0016, a potent and selective inhibitor of 20-HETE synthesizing enzyme. *Br J Pharmacol* **133**:325–329. doi:10.1038/sj.bjp.0704101

Perrenoud Q, Geoffroy H, Gautier B, Rancillac A, Alfonsi F, Tekki-Kessaris N, Rossier J, Vitalis T, Gallopin T. 2012. Characterisation of type I and type II nNOS-expressing interneurons in the barrel cortex of mouse. *Front Neural Circuits* **6**:1–31. doi:10.3389/fncir.2012.00036

Rudolf K, Eberlein W, Engel W, Wieland HA, Willim KD, Entzeroth M, Wienen W, Beck-Sickinger AG, Doods HN. 1994. The first highly potent and selective non-peptide neuropeptide Y Y1 receptor antagonist: BIBP3226. *Eur J Pharmacol* **271**:5–7. doi:10.1016/0014-2999(94)90822-2

Säfholm J, Dahlén SE, Adner M. 2013a. Antagonising EP1 and EP2 receptors reveal that the TP receptor mediates a component of antigen-induced contraction of the guinea pig trachea. *Eur J Pharmacol* **718**:277–282. doi:10.1016/j.ejphar.2013.08.021

Säfholm J, Dahlén SE, Delin I, Maxey K, Stark K, Cardell LO, Adner M. 2013b. PGE2maintains the tone of the guinea pig trachea through a balance between activation of contractile EP1 receptors and relaxant EP2receptors. *Br J Pharmacol* **168**:794–806. doi:10.1111/j.1476-5381.2012.02189.x

Smith CJ, Zhang Y, Koboldt CM, Muhammad J, Zweifel BS, Shaffer A, Talley JJ, Masferrer JL, Seibert K, Isakson PC. 1998. Pharmacological analysis of cyclooxygenase-1 in inflammation. *Proc Natl Acad Sci U S A* **95**:13313–13318. doi:10.1073/pnas.95.22.13313

Sun Q-Q, Baraban SC, Prince DA, Huguenard JR. 2003. Target-Specific Neuropeptide Y-Ergic Synaptic Inhibition and Its Network Consequences within the Mammalian Thalamus.

Tammaro P, Smith AL, Hutchings SR, Smirnov S V. 2004. Pharmacological evidence for a key role of voltage-gated K + channels in the function of rat aortic smooth muscle cells. *Br J Pharmacol* **143**:303–317. doi:10.1038/sj.bjp.0705957
